# A trait-based metric to prioritise the conservation of functionally irreplaceable species at risk of extinction

**DOI:** 10.1101/2024.06.05.597292

**Authors:** Ceri Webster, Joanna Barker, David Curnick, Matthew Gollock, James Hansford, Michael Hoffmann, Thomas E. Martin, Thomas J. Matthews, Nathalie Pettorelli, Joseph A. Tobias, Samuel T. Turvey, Patrick A. Walkden, Jiaqi Wang, Joseph P. Wayman, James Rosindell, Rikki Gumbs

## Abstract

Robust species-level methods for quantifying ecological differences have yet to be incorporated into conservation strategies. Here, we present a conservation prioritisation approach that integrates species trait data and extinction risk to quantify the contribution of individual species to overall functional diversity. The Functionally Irreplaceable with Risk of Extinction (FIRE) metric directs conservation action to species whose extinction is expected to result in significant losses of functional diversity. We applied our framework to sets of species at the global scale. First we assessed the world’s birds, highlighting congruent and divergent priorities identified by trait-based and phylogenetic approaches. Second, we applied FIRE to the world’s sharks, exploring the impact of imputed traits on prioritisation robustness. For birds and sharks, we show that prioritising by functional irreplaceability is an effective strategy to conserve exploited species. The FIRE metric provides a robust tool to facilitate the incorporation of functional diversity into conservation policy and practice, revealing species that may be overlooked by existing approaches.

## 1. Introduction

The biodiversity crisis is driving species declines and losses across all biomes, threatening the functioning of healthy ecosystems and the services they provide^1^. Efforts to mitigate these declines require strategic decision-making and often utilise established metrics of risk or priority, such as conservation status^2^, taxonomic diversity^3^, endemism^4^, climate vulnerability^5,6^ and evolutionary history^7^. Despite increasing interest in the variation of functional traits between species (i.e. functional diversity^8–10^), relatively little progress has been made at incorporating this into conservation decision-making^11^. For example, Goal A of the United Nations Convention on Biological Diversity (CBD) Kunming-Montreal Global Biodiversity Framework (GBF) focuses on maintaining, restoring and enhancing biodiversity to improve functions and services^12^, yet the accompanying monitoring framework contains no means for tracking progress towards recovery of ecological integrity and functional diversity^13^.

Measuring the functional diversity of species assemblages can help to reveal how extinction will impact ecosystem functions and services^14^, providing an estimate of the ecological role or functionality of different species within a given assemblage, and the uniqueness or redundancy of that role in comparison to related or co-occurring taxa. Recent studies suggest that loss of species with functionally unique traits will drive functional homogenisation and erosion of ecosystem processes^15,16^. However, current prioritisation metrics that consider taxonomic or phylogenetic information alone do not adequately capture functional diversity^17,18^.

Advances in metrics that prioritise species for conservation based on phylogenetic diversity (the sum of phylogenetic branch lengths connecting a set of taxa^19^) have produced approaches that are based on the established conceptual framework of averting expected losses of the most imperilled biodiversity^20–24^. For example, the updated Evolutionarily Distinct and Globally Endangered (EDGE) metric, based on the EDGE2 protocol^20^, prioritises species based on the amount of threatened phylogenetic diversity their conservation is expected to secure. In contrast, existing approaches to integrate functional traits into conservation prioritisation metrics have largely been based on the original EDGE metric^7^, which weighted a measure of distinctiveness by an index of extinction risk in an *ad hoc* manner^25^ (though see Pavione^26^). The ‘EcoDGE’^27^ metric is an adaptation of the original EDGE metric that calculates the ‘distinctiveness’ component of the metric using dendrograms representing functional diversity rather than phylogenetic trees. The ‘FUSE’^28^ metric is another adaptation of the original EDGE calculation, whereby an index of priority is created by summing two measures; functional uniqueness^29^ and functional specialisation^29^, both of which are independently weighted by an ordinal ranking of extinction risk. However, growing focus on the maintenance of functionally diverse systems for conservation^8,9^, the increasing availability of trait data across the tree of life^18,30^, and established methods to quantify functional diversity^31^ provide the opportunity to develop a more clearly mechanistic prioritisation approach to guide the conservation of functionally irreplaceable species.

Here, we detail a method that prioritises functionally irreplaceable and threatened species for conservation based on their expected contributions to trait space, given the landscape of extinction risk across the entirety of trait space. We present the Functionally Irreplaceable with Risk of Extinction (FIRE) metric, which builds on previous approaches^26,28,32^ to quantify the expected loss of functional diversity that can be averted through species-based conservation action. We then provide two case studies illustrating the utility of our new approach. The first applies FIRE to the world’s birds, using a comprehensive species-level trait dataset for over 10,000 species^33^ to highlight unique and convergent conservation priorities for maintaining avian functional and phylogenetic diversity. For the second, we apply FIRE to 557 of the world’s shark species, a smaller taxonomic group with more patchy data, allowing us to explore how imputation of missing data influences the setting of conservation priorities using the FIRE approach. We then use functional trait data for birds and sharks to examine whether species threatened by hunting, collecting, and fishing (i.e. exploitation) are particularly functionally or evolutionarily irreplaceable, and explore how conservation strategies to prioritise irreplaceable species perform at safeguarding diversity at risk from this threat. FIRE builds on existing prioritisation frameworks to provide functional indices to inform conservation and policy.

## 2. Results

### 2.1 Functionally Irreplaceable with Risk of Extinction (FIRE) metric

The FIRE metric aims to identify and prioritise threatened species whose extinction is expected to represent disproportionately large losses of unique functional trait space. FIRE estimates the functional irreplaceability of a species by calculating its expected future unique contribution to overall trait space given the extinction risk and functionality of all other species in the same trait space. The trait space from which FIRE is calculated requires independent trait axes to quantify functional distances between species (Fig. 1, see Methods). In the case of continuous traits, including morphometric data, these independent axes can be generated using ordination methods (PCA or PCoA, depending on data type). Trait probability densities (TPDs) are then used to estimate a probabilistic trait space based on the results of the ordination (PCA or PCoA)^34^. The expected loss of overall trait space is then calculated using a large number (n ≥ 1000) of extinction iterations where, for each iteration, species are removed from trait space with probabilities derived from their IUCN Red List categories (Fig. 1, see Methods). For each species, a distribution of functional irreplaceability scores is then calculated by quantifying the amount of trait space expected to be uniquely occupied by the species if it were to avoid extinction in each of the *n* extinction iterations. The FIRE of a species is then calculated as the product of its functional irreplaceability and probability of extinction across the distribution of *n* iterations, and represents the expected loss of trait space that could be averted through the conservation of each species (i.e., if the species does not go extinct) (Fig. 1). To facilitate comparability between functional irreplaceability scores calculated from different trait spaces (for a fixed taxonomic group), we present the scores as the percentage of total trait space for which we expect a species to be responsible in the future.

**Fig. 1.**
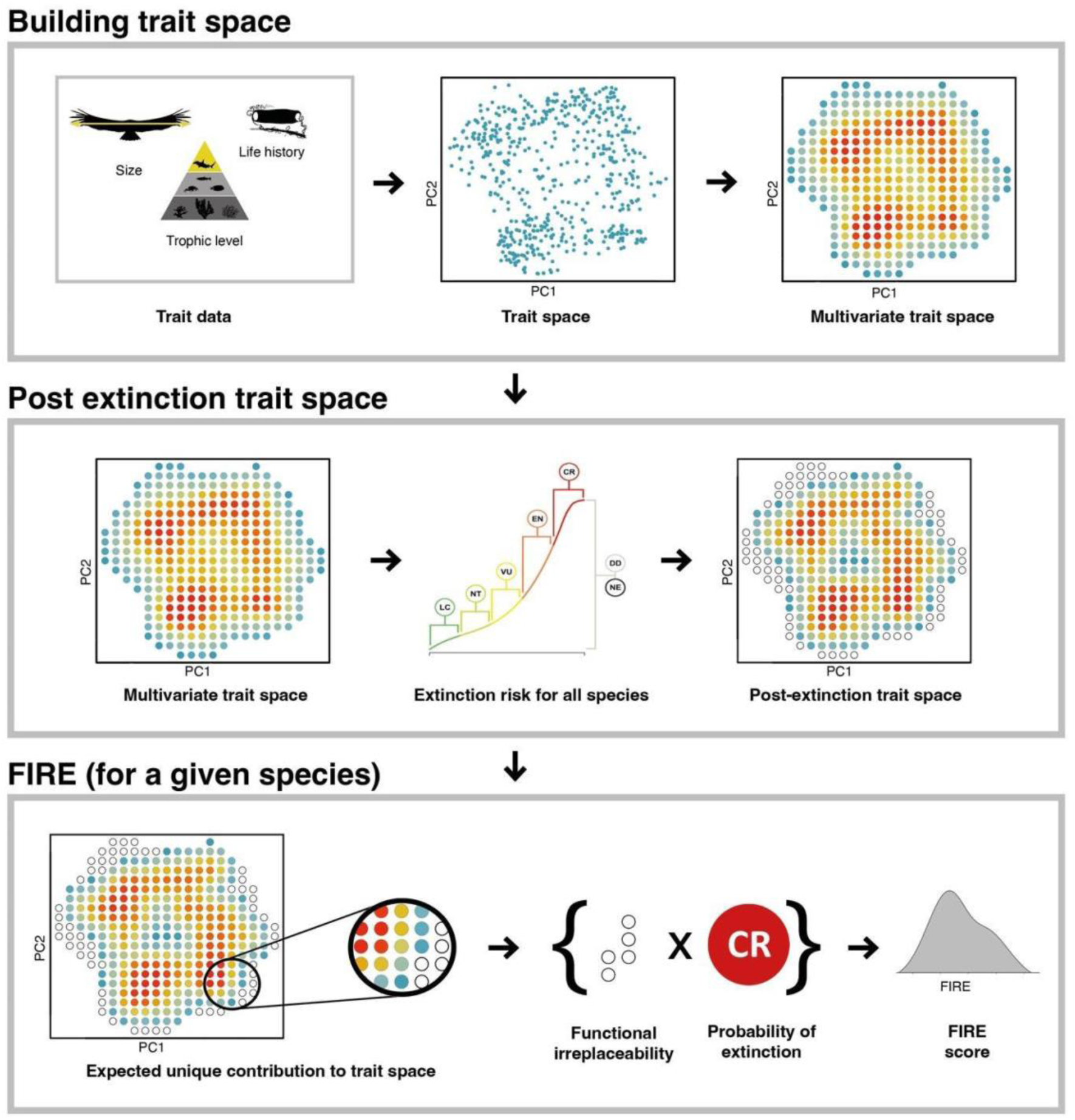
| Calculating the Functionally Irreplaceable with Risk of Extinction (FIRE) metric. Top panel: Constructing a multivariate trait space requires trait data to quantify functional distances between species. Probabilistic trait space is estimated from the functional distances between species and an estimated variability using the TPD package (v1.1.0)^34^. Colours in trait space indicate high (red) or low (blue) probability of occupancy. Middle panel: Post-extinction trait space is calculated by removing species predicted to go extinct in a given iteration following an extinction scenario, whereby species become extinct or survive based on their probability of extinction. White circles represent areas of trait space no longer occupied by species following extinction. This step is repeated across 1000 iterations to derive a distribution of uncertainty in extinction scenarios. IUCN Red List categories are converted to a continuous distribution of extinction probabilities, following Gumbs et al. 2023^20^ where the median of each Red List category was defined as; Critically Endangered = 0.97, Endangered = 0.485, Vulnerable =0.2425, Near Threatened = 0.12125, and Least Concern = 0.060625. Species probability of extinction is randomly selected from the distribution of extinction probabilities defined by their IUCN Red List categories. Species that were not assessed or had IUCN Red List category listed as Data Deficient, were randomly sampled from the entire distribution of extinction probabilities. Bottom panel: The functional irreplaceability of a species is calculated as the distinct contribution of that species (which is immune to extinction during the calculation of its own functional irreplaceability score) to post-extinction trait space. In the example shown there are five pixels in discretised trait space that will in future only be covered by the focal species, if the focal species is protected from extinction. The FIRE value of a species is then calculated as the product of its functional irreplaceability and probability of extinction. The calculation is repeated 1000 times to capture the distribution of outcomes from the probability of extinction. Image credits given in supplementary material.

### 2.2 Case study 1: FIRE for the world’s birds

Birds represent an ideal taxonomic group for testing the FIRE metric, given their long-standing role in conservation research^35^, the availability of comprehensive, high-resolution trait data^33,36^, and a robust literature base linking traits to ecological functions^37^.

#### 2.2.1 Trait space summary

To quantify the trait space of the world’s birds (11,005 species), we selected eight morphological traits that have been shown to provide accurate information on the functional role and trophic status of birds at the global scale^33^: total beak length (from the tip to the skull), beak length to the nares, beak width and depth, wing length, secondary length (length from the carpal joint to the first of the wings secondary feathers), tail length and tarsus length. We found consistently weak correlations between morphological traits and extinction risk. Tarsus length (ρ = 0.160, p < 0.001), beak depth (ρ = 0.157, p < 0.001), and secondary length (ρ = 0.154, p < 0.001) showed the strongest associations, though effect sizes remained low. Other traits such as beak length (culmen: ρ = 0.155; nares: ρ = 0.147), wing length (ρ = 0.149), and beak width (ρ = 0.133) also showed significant but weak positive associations (all p < 0.001). Tail length had the weakest relationship with extinction risk (ρ = 0.110, p < 0.001). Overall, the low ρ values across traits suggest limited predictive value of individual morphological variables for extinction vulnerability (Fig. S1).

We undertook a Principal Components Analysis (PCA) using these eight traits and extracted the first three axes, which together represent 92% of the total variation. PCA axis 1 (PC1; 76% of total variation) represents an overall size axis, PC2 (10%) represents a trade-off between beak length vs. tarsus, secondary and tail length, and PC3 (6%) represents a trade-off between beak length vs. beak width and depth. We re-ran the analyses including body mass as an additional trait. The functional irreplaceability (r = 0.96, p < 0.001) and FIRE (r = 0.97, p < 0.001) values were highly correlated between the two approaches and we only discuss the results excluding body mass from herein. We used TPDs to account for the probabilistic nature of species traits^38^ (see Methods).

To estimate the predicted loss of trait space in the future, we projected species risk of extinction onto global trait space (see Methods). Mapping extinction risk onto trait space revealed marked variation in vulnerability (Fig. 2). We show that extinction in trait space is likely to occur as a loss of distinct clusters rather than as a uniform contraction around the periphery.

**Fig. 2.**
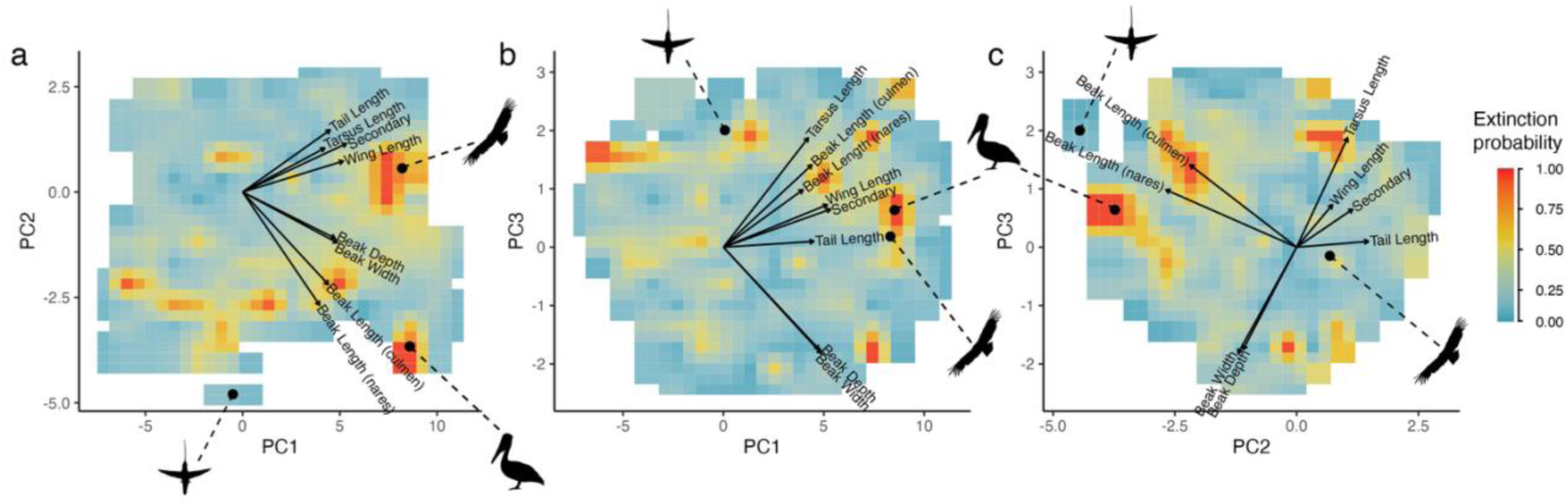
| Extinction risk of avian trait space. **a-c**, Trait spaces defined by three principal component axes (PC1, PC2 and PC3) to show; the direction and weighting of the considered traits (black arrows), and projections of the predicted erosion of trait space based on IUCN Red List assessments^2^. Light blue indicates low probability of extinction and red indicates high probability of extinction. Dashed lines show examples of species position in trait space, including California condor (*Gymnogyps californianus*), Brown pelican (*Pelecanus occidentalis*) and Sword-billed hummingbird (*Ensifera enifera*). Traits include; total beak length (from the tip to the skull), beak length to the nares, beak width and depth, wing length, secondary length (length from the carpal joint to the first of the wings secondary feathers), tail length and tarsus length. Image credits given in supplementary material.

#### 2.2.2 Avian functional irreplaceability

The world’s most functionally irreplaceable birds include several species widely considered to be morphologically and ecologically unique, such as the Shoebill (*Balaeniceps rex*), Great Hornbill (*Buceros bicornis*), Sunbittern (*Eurypyga helias*), kiwis (*Apteryx* spp.), and Secretarybird (*Sagittarius serpentarius*; Fig. 3a). Other unique species are more obscure, including the Coppery Thorntail (*Discosura letitiae*; DD on Red List) which is only known by a single museum specimen of unknown provenance. Overall, there was a trend for hummingbird species to be relatively highly ranked in the list of bird functional irreplaceability values (discussed in more detail in Text S1). Conversely, the least functionally irreplaceable species tend to be small passerines (e.g. warblers, tits; see full list in Data S1). Eleven bird species do not overlap with any other species in trait space and thus receive maximum irreplaceability scores (Data S1).

**Fig. 3.**
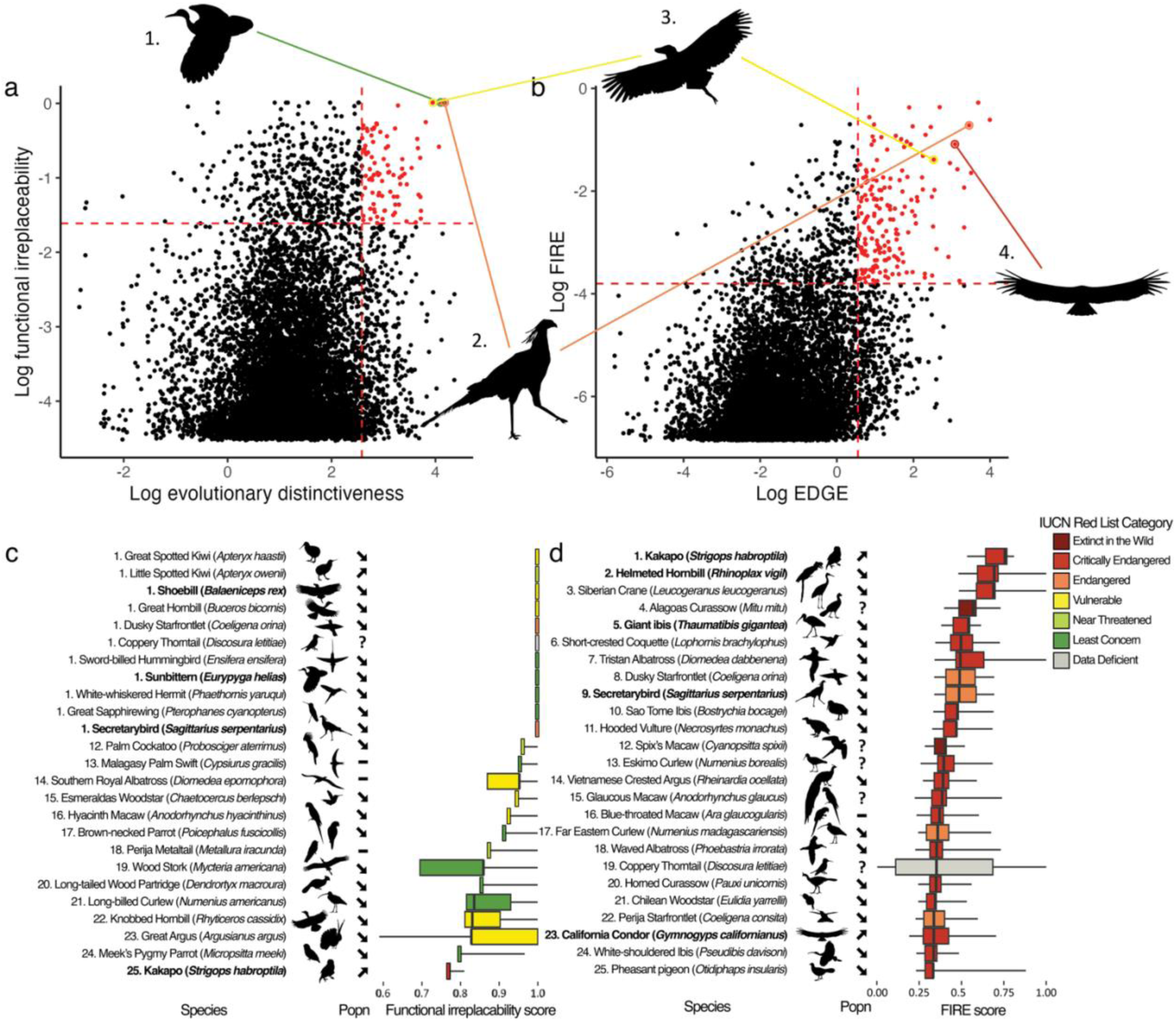
| Functionally irreplaceable birds. Correlations between (a) evolutionary distinctiveness and functional irreplaceability and (b) Evolutionarily Distinct and Globally Endangered (EDGE) and Functionally Irreplaceable with Risk of Extinction (FIRE) scores. Dotted lines represent the upper 5% of scores for each metric. Illustrations represent examples of species that fall within the top 5% of (c) evolutionary distinctiveness and functional irreplaceability (d) EDGE and FIRE. Bird species: 1. Sunbittern (*Eurypyga helias*), 2. Secretarybird (*Sagittarius serpentarius*), 3. Shoebill (*Balaeniceps rex*), 4. California Condor (*Gymnogyps californianus*). The top 25 highest ranked (c) functionally irreplaceable and (d) FIRE bird species and numbers represent the (c) functional irreplaceability and (d) FIRE rank of each species, where species with the same median value have the same rank. Boxplots represent the range, interquartile range and median of (c) functional irreplaceability and (d) FIRE scores calculated over 1000 iterations. IUCN Red List population trends (Popn) are denoted with arrows to represent increasing or decreasing population trends, a dash to represent stable population trends, and a question mark to represent unknown population trends. Species in bold text fall in the top 5% of both (c) functional irreplaceability and evolutionary distinctiveness scores and (d) FIRE and EDGE scores. Colours represent IUCN Red List categories (red = Critically Endangered, orange = Endangered, yellow = Vulnerable, light green = Near Threatened, green = Least Concern, grey = Data Deficient). Image credits given in supplementary material.

Threatened birds (i.e. Vulnerable [VU], Endangered [EN], and Critically Endangered [CR] on the Red List) are significantly more functionally irreplaceable than non-threatened (i.e. Least Concern [LC] and Near Threatened [NT]) birds (mean = 0.065 vs. 0.035, Welch’s t-test: df = 1486.7, p < 0.001), corroborating earlier work highlighting this relationship^15,39,40^. Of the top 5% most functionally irreplaceable species (N = 551), 25.4% are threatened with extinction, compared with 11.7% of all birds^2^. The highest priority FIRE species (i.e. highly functionally irreplaceable species threatened with extinction) include the Extinct in the Wild Alagoas Curassow (*Mitu mitu*), and the Critically Endangered Helmeted Hornbill (*Rhinoplax vigil*) and Tristan Albatross (*Diomedea dabbenena*), alongside species such as the Secretarybird (Endangered), Shoebill (Vulnerable), and Kākāpō (*Strigops habroptila*; Critically Endangered; Fig. 3d; see full list in Data S1).

#### 2.2.3 Unique and convergent FIRE priorities

To compare convergent and unique conservation priorities between FIRE and the phylogenetically-informed EDGE approach, we matched the bird species in our FIRE dataset with those with available EDGE data (N = 10971)^41^. There is a weak but significant positive correlation between functional irreplaceability and evolutionary distinctiveness for the world’s birds (ρ = 0.1, df = 10684, p < 0.001; Fig. 3a), though species that comprise monotypic families have significantly higher functional irreplaceability than birds in general (mean = 0.15 vs. 0.04, Welch’s t-test: df = 33.021, p = 0.03). We identified species within the highest 5% of FIRE scores, in a threatened or Extinct in the Wild Red List category, and with available EDGE scores, as our set of priority birds (N = 316). Of these priority 5% species, 165 are also in the top 5% of EDGE scores, and include species such as the Helmeted Hornbill, Secretarybird, Shoebill, Andean Condor (*Vultur gryphus*), and New Caledonian Owlet-nightjar (*Aegotheles savesi*). We highlight these species as amongst the most important for conservation, given their particularly high contribution to irreplaceable functional and phylogenetic diversity globally (Fig. 3b, full list in Data S1).

Conversely, 102 (32.3%) of the top 5% FIRE species do not qualify as priority EDGE species (i.e. above median EDGE with 95% confidence and threatened with extinction^20^), and thus are not currently captured by phylogenetically-informed prioritisation. Further, 71 of the top 5% FIRE species do not trigger any Key Biodiversity Areas (KBAs), and 45 (14.2%) are underrepresented in the global KBA network (i.e. <8% of species area of habitat (AOH) covered by KBAs; Lansley et al. 2025). While many of these species are nevertheless the subject of targeted conservation actions (e.g. the Okarito Kiwi, *Apteryx rowi*^42^), additional conservation efforts are likely required to promote and conserve some of the world’s most functionally irreplaceable birds. For example, FIRE scores tend to shift the focus of conservation attention from less distinctive lineages towards maintaining interconnected populations of widespread species with important ecological roles, such as Kori Bustard *Ardeotis kori* and Greater Rhea *Rhea americana*.

### 2.3 Case study 2: FIRE prioritisation with incomplete data

Given the scale and urgency of the biodiversity crisis, it is impractical to limit conservation research and efforts to only a select few taxonomic groups or regions with complete data ^43–45^. Major advances have been made to facilitate the inclusion of species lacking data into assessments of extinction risk^46,47^, conservation priorities^20,48^, and functional diversity^49^. We designed the FIRE metric to permit the inclusion of species lacking extinction risk data (Fig. 1). In addition, the metric can be calculated using imputed trait data^84^. To illustrate the calculation of FIRE for a group with good – but incomplete – trait data coverage, we applied the metric to the world’s sharks (Selachimorpha). A substantial amount of trait and extinction risk data are now available for the world’s sharks, a group of high ecological importance and significant conservation concern^28,50^. Given their elevated extinction risk and functional roles in marine ecosystems, sharks have been the subject of previous trait-based prioritisation efforts based on imputed data^32^.

#### 2.3.1 Imputing trait space

To quantify the trait space occupied by the world’s sharks, we selected eight traits that characterise a variety of functional guilds (maximum body length, maximum depth in the water, trophic level, reproductive guild, habitat preference, growth ratio, and the first two principal component axes from an analysis of body shape; see Methods) for 557 species (>99% of all sharks). Taxonomic coverage of these eight traits was incomplete but high overall (maximum body length: 96.1%, maximum depth: 87.6%, trophic level: 49.9%, body shape variation PC1: 91.9%, body shape variation PC2: 91.9%, growth ratio: 61.4%, reproductive mode: 84.2%, habitat preference: 100%; Table S1).

We used random forest models (missForest)^51^ to impute missing trait values and attain 100% coverage for all traits. This method uses a random forest trained on the observed values of the trait matrix to impute missing values. MissForest makes no prior assumptions about the distributions of variables, does not incorporate phylogenetic relationships and can be used to predict both continuous and categorical data^51^. Additionally, numerous studies have found a high degree of accuracy when imputing traits using random forest methods such as missForest^49,52,53^. Following imputation of all traits, we found generally weak or inconsistent associations between traits and extinction risk. Shape PC1 and body length showed the strongest relationships (ρ = 0.41 and ρ = 0.44, respectively; both p < 0.001). Maximum depth (ρ = -0.20, p < 0.001) and growth ratio (ρ = 0.16, p < 0.001) showed modest effects. Traits such as trophic level (ρ = 0.06, p = 0.15) and shape PC2 (ρ = -0.09, p = 0.05) were weak predictors, with minimal explanatory power (Fig. S2). Categorical traits showed significant differences in extinction score among groups for both habitat preference (Kruskal-Wallis χ² = 53.54, df = 7, p < 0.001) and reproductive guild (Kruskal-Wallis χ² = 56.23, df = 3, p < 0.001). However, these differences were not clearly directional and did not indicate a consistent trait gradient with increasing extinction risk (Fig. S2).

We defined trait space occupied by extant sharks by condensing the dissimilarity distance matrix of compiled traits using a principal coordinate analysis (PCoA), and extracting the first four axes, which together represent 45.28% of total variation. To assess how imputation was distributed across trait space, we calculated the proportion of imputed data for each species and projected this onto the first two PCoA axes (Fig. S3). We then summarised imputation density across trait space by calculating the average proportion of imputed data along both axes. This analysis revealed that imputation was relatively evenly distributed, with no strong clustering of highly imputed species in any particular region of trait space. Average imputation values along PC1 ranged from 0 to 0.26 (median: 0.18), and along PC2 from 0.05 to 0.32 (median: 0.15; Fig. S3). Species with a high proportion of imputed data (more than 50% of data imputed) did not have high functionally irreplaceability scores (Fig. 4a).

**Fig. 4.**
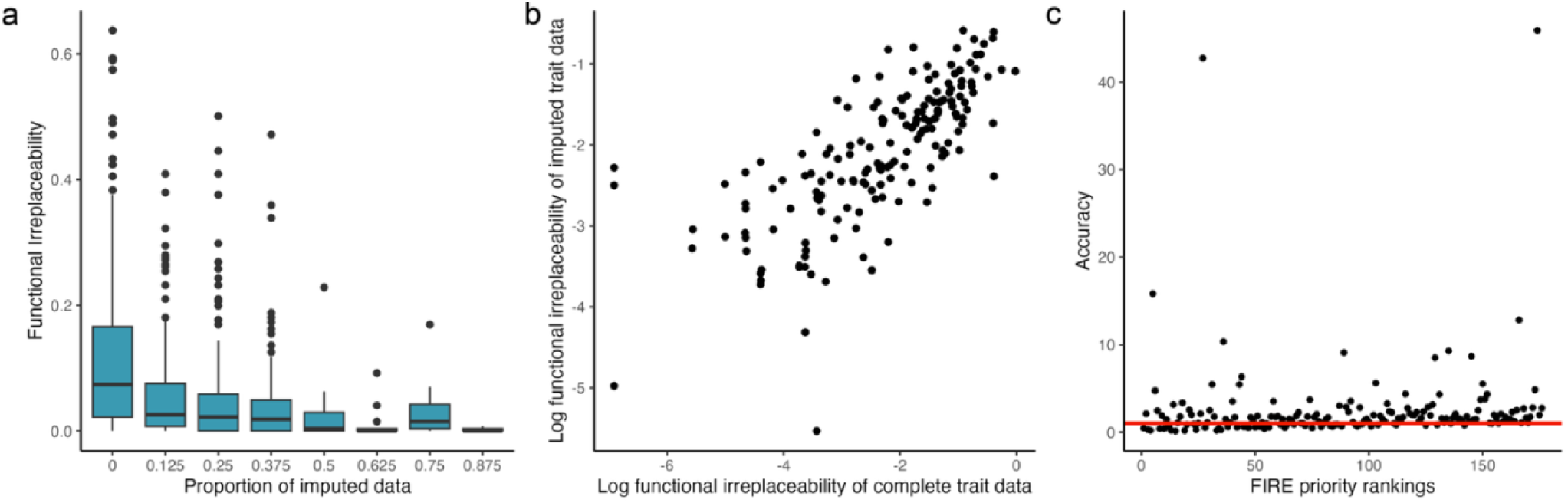
| Performance of functional irreplaceability estimations using imputed data for shark species. (a) The proportion of trait data imputed for species in our complete shark dataset (N = 557) and their functional irreplaceability scores. (b) The correlation between functional irreplaceability scores of the gapless dataset (no missing trait data; and a simulated imputed dataset (N = 176). (c) The change in FIRE rankings between the full FIRE dataset and the gapless dataset (Accuracy) for species with full trait data availability (N = 176).

To test the validity of our trait space derived from real and imputed data, we calculated functional irreplaceability scores from a trait space constructed only with species with complete trait data (‘gapless dataset’; N = 176). We then generated artificially incomplete data by randomly removing trait values at the observed proportion of incompleteness, then imputing them back. We found a strong correlation between the median functional irreplaceability scores from the gapless dataset and those from the imputed datasets (ρ = 0.78, p < 0.001; see Methods and Fig. 4b). The change in FIRE rankings of the 176 species in the gapless dataset, when compared with their rankings in the full (imputed) dataset, was consistent irrespective of rank (ρ = 0.08, df = 175, p = 0.3; Fig. 4c), indicating that the inclusion of species with incomplete trait data did not affect priority rankings in a biased manner. Indeed, 27 of 34 species (79%) remained in the highest 20% of rankings, and 13 of 18 (72%) remained in the top 10% across both datasets. However, the robustness of these results is likely to be sensitive to the traits selected and the imputation method used, particularly if trait gaps are non-randomly distributed or if certain traits disproportionately influence ordination and irreplaceability scores^49,53^.

#### 2.3.3 Priority FIRE species

The species with the highest FIRE scores are the Sand Tiger Shark (*Carcharias taurus*; representing 0.44% avertable expected loss of trait space), Scalloped Hammerhead (*Sphyrna lewini*; representing 0.33% avertable expected loss) and Basking Shark (0.31%; Fig. 5). Shark orders Echinorhiniformes and Lamniformes have the highest average FIRE scores of 0.08% (sd = 0.02) and 0.07% (sd = 0.12), respectively (more information on functional irreplaceability and extinction risk in Text S2). We identified 76 shark species as priority FIRE species, defined as threatened sharks for which we have 95% certainty that they are above median FIRE (mirroring the approach to identifying priority EDGE species^20^; full list in Data S2). All of the top 76 priority FIRE sharks and 67.9% of the 215 priority functionally irreplaceable sharks have declining or unknown population trends, based on Red List data^2^. One Critically Endangered species in the top 28 FIRE sharks, the Lost Shark (*Carcharhinus obsoletus*), is Possibly Extinct, and has not been recorded since 1934^54^.

**Fig. 5.**
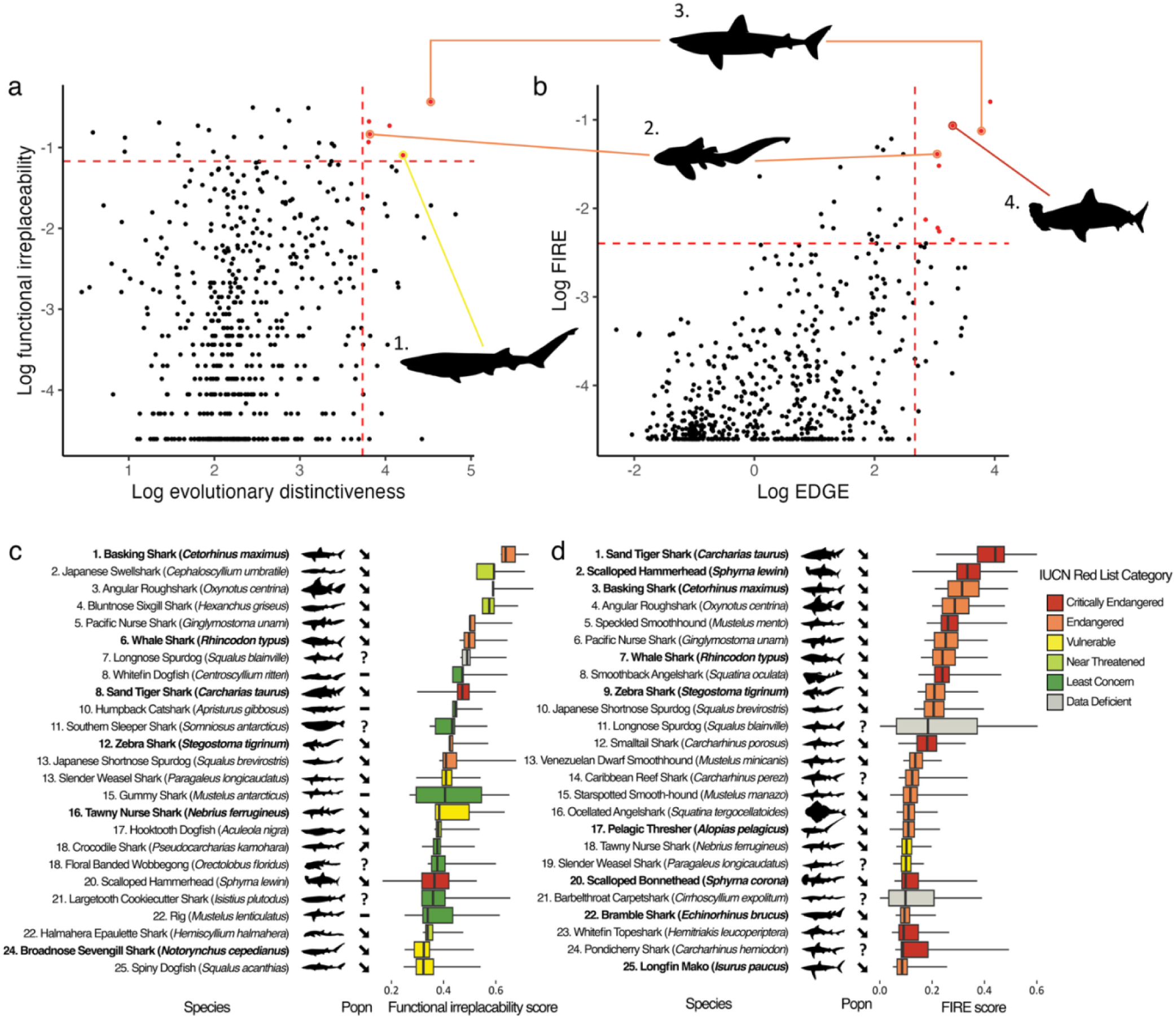
| Functionally irreplaceable sharks. Correlations between (a) evolutionary distinctiveness and functional irreplaceability and (b) Evolutionarily Distinct and Globally Endangered (EDGE) and Functionally Irreplaceable with Risk of Extinction (FIRE) scores. Dotted lines represent the upper 5% of scores for each metric. Silhouettes represent examples of species that fall within the top 5% of (c) evolutionary distinctiveness and functional irreplaceability (d) EDGE and FIRE. Shark species: 1. Broadnose Sevengill Shark (*Notorynchus cepedianus*), 2. Zebra Shark (*Stegostoma tigrinum*), 3. Basking Shark (*Cetorhinus maximus*), 4. Scalloped Hammerhead (*Sphyrna lewini*). The top 25 highest ranked (c) functionally irreplaceable and (d) FIRE shark species. Numbers represent the (c) functional irreplaceability and (d) FIRE rank of each species, where species with the same median value have the same rank. Boxplots represent the range, interquartile range and median of (c) functional irreplaceability and (d) FIRE scores calculated over 1000 iterations. IUCN Red List population trends (Popn) are denoted with arrows to represent increasing or decreasing population trends, a dash to represent stable population trends, and a question mark to represent unknown population trends. Species in bold text fall in the top 25 of both (c) functional irreplaceability and evolutionary distinctiveness scores and (d) FIRE and EDGE scores. Colours represent IUCN Red List categories (red = Critically Endangered, orange = Endangered, yellow = Vulnerable, light green = Near Threatened, green = Least Concern, grey = Data Deficient). Silhouette credit available in supplementary material.

#### 2.3.4 Relationship between FIRE and EDGE

We applied the EDGE2 protocol^20^ to calculate evolutionary distinctiveness and EDGE scores for sharks, and compared these with functional irreplaceability and FIRE. Functional irreplaceability and evolutionary distinctiveness showed a weak but significant positive correlation (Pearson’s r = 0.18, p < 0.001; Fig. 5a). Six species ranked in the top 5% for both metrics, while three of the top 5% of functionally irreplaceable species, including the Southern Sleeper Shark (*Somniosus antarcticus*) and Gummy Shark (*Mustelus antarcticus*), were among the bottom 5% for evolutionary distinctiveness. Ten species ranked in the top 5% for both FIRE and EDGE scores (Fig. 5b), with the Sand Tiger Shark (rank 1 for both), Basking Shark (FIRE rank 3, EDGE rank 2), and Scalloped Hammerhead (FIRE rank 2, EDGE rank 9) ranking highest across both frameworks.

### 2.4 Case study 3: Functional irreplaceability and exploitation

To determine whether there is a relationship between the different distinctiveness measures (evolutionary distinctiveness and functional irreplaceability) and species targeted for human exploitation, we selected sets of species of increasing richness based on functional irreplaceability and based on evolutionary distinctiveness. Specifically, species with higher scores in a given metric had a proportionally greater probability of being selected in a set, we also used randomly selected sets for comparison (see Methods). There are 468 shark species (84.0% of total) with biological resource use (BRU; i.e. intentional and unintentional fishing) listed as a threat on the IUCN Red List^2^. For 168 species (30.2% of total), BRU is listed as intentional, and for 300 (53.9% of total), BRU is listed as unintentional. There are 1256 bird species with intentional BRU (i.e. hunting and collecting) listed as a threat, with an additional 125 species threatened by unintentional BRU^2^ (see Methods).

For birds, selecting species based on functional irreplaceability highlighted more intentionally exploited species than evolutionary distinctiveness at all sample sizes (Fig. 6a; Table S2). For sharks, functional irreplaceability again highlights the greatest number of intentionally exploited species (Fig. 6b; Table S2). Selecting shark species based on evolutionary distinctiveness performs significantly better than random at highlighting intentionally exploited species (Fig. 6b; ANOVA with Tukey’s test all p < 0.001 compared with random; all pairwise comparison results available in Table S2).

**Fig. 6.**
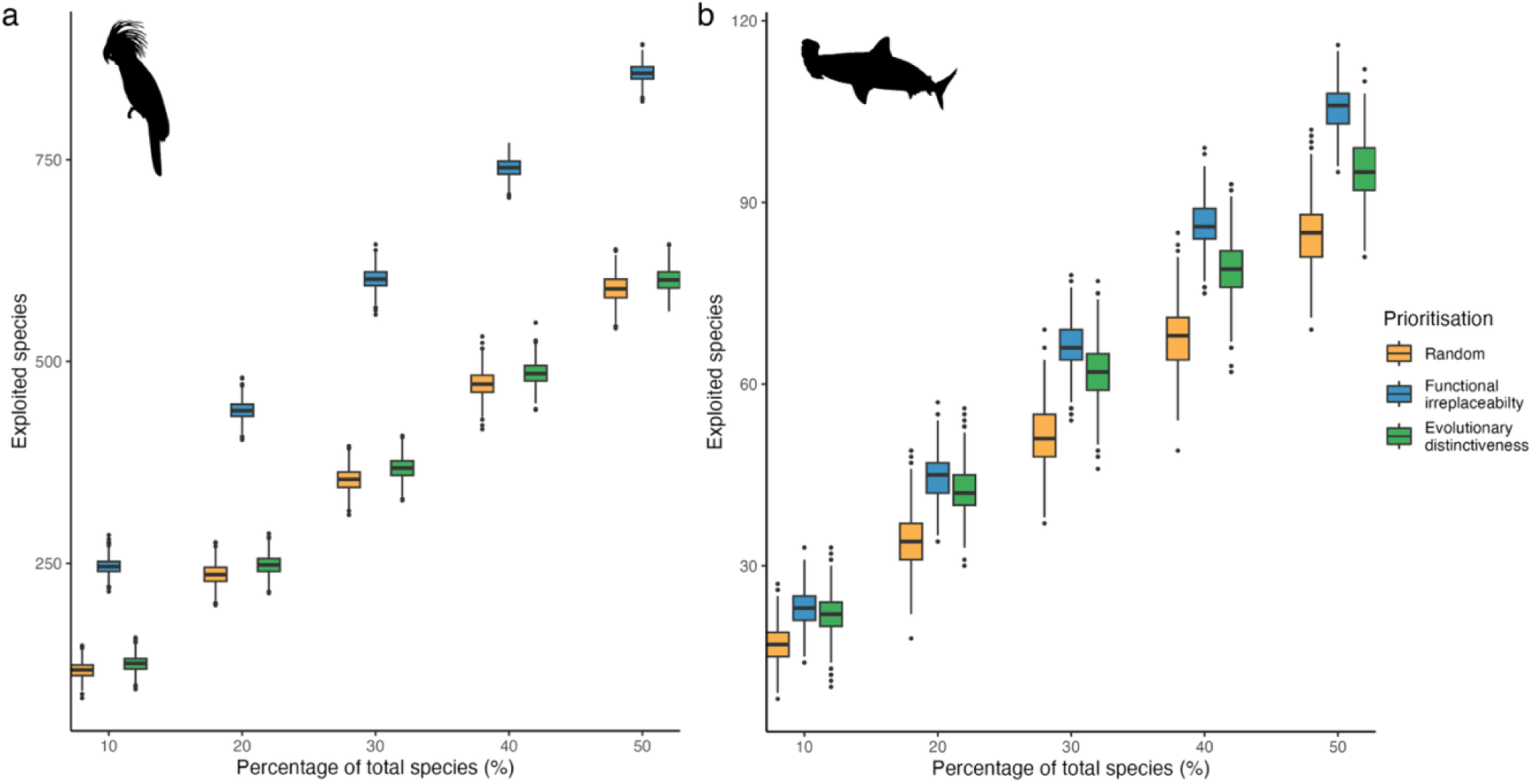
| The relationship between intentional exploitation of species and their Functional Irreplaceability and Evolutionary Distinctiveness. The number of intentionally exploited a) bird and b) shark species highlighted when 10%-50% of species are selected for conservation based on weighted random draws from different distributions corresponding to different conservation strategies: uniform (fully random; orange), weighted by functional irreplaceability (blue) or weighted by evolutionary distinctiveness (green). Boxplots represent median value (solid line), 25^th^ and 75^th^ percentiles (box edges), and 5^th^ and 95^th^ percentiles (whiskers) from 1000 iterations. Silhouette credit available in supplementary material.

For both birds and sharks, species with intentional BRU listed as a threat had significantly higher functional irreplaceability compared to species where intentional BRU is not listed as a threat (sharks: mean = 0.10 vs. 0.06, df = 249.1, p < 0.001; birds: mean = 0.09 vs. 0.03, df = 1227.5, p < 0.001) and evolutionary distinctiveness (sharks: mean = 18.9 vs. 12.4, df = 225.1, p < 0.001; birds: mean = 5.3 vs. 4.7, df = 1227.5, p = 0.001).

## 3. Discussion

The FIRE metric offers a novel approach to identify priority species whose conservation is expected to avert the greatest losses of irreplaceable functional diversity. Given the wealth of evidence linking functional diversity to the provision of ecosystem processes and services, it is vital to prevent further erosion of functional trait space^55–57^. By applying FIRE to global-scale datasets of birds and sharks, we illustrate the metric’s ability to identify priorities for conservation for different sets of taxa with contrasting levels of species richness and data coverage. Our results highlight numerous species of both birds and sharks that are not currently identified as priorities by phylogenetically-informed approaches, alongside threatened species that are both phylogenetically and functionally irreplaceable. Further, our results indicate that functionally irreplaceable species are disproportionately threatened by exploitation (i.e. hunting, collecting, fishing, according to IUCN Red List data).

The FIRE metric, which measures the expected contribution of species to overall trait space, builds on approaches such as EcoDGE^27^ and FUSE^28^ to integrate functional diversity into an expected loss framework. As FIRE incorporates complementarity to quantify the risk to each point in trait space due to the extinction risk of the species present, the approach also presents further opportunities beyond species prioritisation. Our methods can be used to measure the expected erosion of trait space due to extinctions, and to quantify variation in the probability of occupancy in regions of trait space as extinction risk changes over time. Through these advances, FIRE can potentially underpin the development of a coherent suite of metrics to monitor risk to functional diversity, and guide conservation efforts, as has been achieved with the EDGE and Phylogenetic Diversity indicators adopted by the GBF^58^. We calculated FIRE for birds and sharks at the global scale. However, FIRE can also be calculated at regional, national or community levels. These complementary levels of analysis would allow us to identify species whose function within specific ecosystems or localities is expected to be distinct and at risk of being lost^59,60^. However, the highlighted species in this context may not be irreplaceable as they are with global scale analyses, because they could be (re)introduced from other localities outside the focal area. Such assessments nevertheless have strong potential to inform local, regional and global conservation planning and policy.

The trait space underpinning FIRE calculations is generated using TPDs, as we wanted to account for the probabilistic nature of species traits and consider intraspecific trait variability when describing functional space^38^. TPDs measure the probability of occupancy across trait space, and FIRE is designed to be calculated with either a binary or probabilistic TPD trait space (see Methods). We used the probabilistic approach to calculate FIRE for birds, given the large number of species and thus high potential for many species to occupy areas of trait space fully occupied by multiple other species, leading to such species having a functional irreplaceability and FIRE score of zero. For our shark case study, which comprised ∼6% of the richness of birds, we illustrated the binary approach to the occupation of trait space by a given species; a species occupies a region of trait space or it does not. As TPDs use kernels to represent the probabilistic distribution of species in trait space, additional work to explore the sensitivity of these approaches to kernel parameters is needed. Future calculations of FIRE may incorporate intraspecific variation to quantitatively parameterise kernel size and occupancy probability across trait space.

Though currently formulated based on the use of TPDs, FIRE in its broadest terms – i.e the amount of expected functional diversity loss that can be averted through conservation action on a single species – can in principle utilise any approach to measuring functional richness (e.g., dendrograms). We simply apply extinction to the set of species and compare total diversity (according to our chosen measure) of a case where our focal species is permitted to go extinct with a case where our focal species is protected. As with all functional diversity analyses, FIRE is sensitive to methodological choices, including the metric used to quantify functional diversity, the traits selected to represent ecological functions, and the number of PCA / PCoA axes retained. Consequently, any given FIRE priority list should be viewed as contingent on these choices, rather than as a single fixed or definitive ranking.

The growing interest in quantifying functional diversity for conservation purposes^61^ has led to an increase in studies examining the patterns and processes of functional diversity and the impacts of biodiversity loss^62^. However, persistent knowledge gaps and limited trait data continue to constrain functional diversity research, often restricting analyses to specific taxonomic groups and geographic regions^17^. As a result, most functional diversity studies have focused on well-documented groups such as plants and birds^33,63^, potentially limiting the generality of their conclusions. This taxonomic and geographic bias underscores the need to broaden the scope of functional diversity assessments. Recent advances in the collection and availability of trait data across a wider range of taxa^64–66^ and the development of methods for imputing missing data^51,67,68^ present new opportunities to do so. In this study, we demonstrate that incomplete datasets can still yield meaningful insights into functional diversity. By calculating FIRE from an incomplete shark trait dataset using imputation, we show that it is possible to assess functional diversity more broadly, even in the face of substantial data gaps. However, reliance on single imputation may overlook the uncertainty in missing data^69^. Multiple-imputation approaches offer a promising avenue for future work, as they better capture this uncertainty^69^.

As trait data availability increases across taxonomic groups^33,64,65,70–72^, the number and types of traits used to inform trait space will vary and will require careful consideration. For example, an important aspect of shark life history that we were unable to include due to a lack of trait availability is changes through different life stages^73^. Many species will display changes in functional and behavioural features, such as shifts in trophic position and foraging location^74^ throughout their lifetime, and future work should aim to incorporate these and other trait dynamics to understand the relationship between ecological function and extinction risk more comprehensively.

The moderate correlation between FIRE and EDGE for both birds and sharks is to be expected due to the incorporation of extinction risk in both metrics. However, there is only a weak correlation between functional irreplaceability and evolutionary distinctiveness for both birds and sharks - a finding that echoes recent studies that challenge the longstanding assumption that phylogenetic diversity should be employed as a surrogate for functional diversity^19,75^. This disparity is expected because of widespread convergent and divergent evolution, which weakens the link between phylogeny and function^37^. Indeed, there is growing evidence that species traits, rather than phylogeny, provide the most accurate index of ecological function in a range of taxonomic groups^37^. Our results thus support the argument that phylogenetic diversity and functional diversity should be considered as two interlinked yet distinct dimensions of biodiversity, both of which are vital components of conservation strategies to preserve ecosystems^56,76,77^ and promote human wellbeing^78,79^.

Indeed, our work further highlights the complex and non-linear relationship between major adaptive change and evolutionary time^80–82^. For sharks, three species were in the top 5% of functional irreplaceability but the bottom 5% of evolutionary distinct species. Exploring the distribution of these species in trait space highlights that there are multiple clusters that are occupied by many species. However, though these three species were clustered with their congeners, they occupy areas on the periphery of their respective genus clusters. Their unique trait combinations cause these species to occupy low density areas, in at least one dimension of trait space, which is not occupied by many other species (Fig. S4). It is well established that certain traits, such as body size and generation length^83^, are associated with increased extinction risk, and it is to be expected that regions of trait space dominated by such traits will be inherently vulnerable^15,16,84^. Further exploration is needed to understand how key traits are distributed across the multiple axes of trait space produced by TPDs. Where intrinsically sensitive traits cluster in sparsely occupied regions of trait space, there is the potential for cascading losses due to functionally clumped extinction risk^15,16^.

Many of the highest-ranking FIRE species have been identified as priorities by earlier approaches using different trait data^32^, calculations of trait space^18,78^, and treatment of extinction risk^84^, supporting the robustness of our identified priorities. While there is moderate congruence in the priority species identified by FIRE and EDGE, there are high-ranking FIRE species that could be overlooked if only focusing on high-ranking EDGE species, such as the Slender-billed Vulture (*Gyps tenuirostris*), Sira Curassow (*Pauxi koepckeae*), Angular Roughshark (*Oxynotus centrina*) and the Pacific Nurse Shark (*Ginglymostoma unami*).

Indeed, prioritising any single biodiversity facet alone risks overlooking species that may significantly contribute to other ecological or phylogenetic aspects of biodiversity. We therefore highlighted species that are in the top 5% of both FIRE and EDGE scores as species of particular interest whose conservation would represent large conservation gains across multiple dimensions of biodiversity (Fig. 3 and 5). These include many iconic threatened bird species, such as the Helmeted Hornbill, Shoebill and Kākāpō, along with high-profile shark species, including Scalloped Hammerhead, Pelagic Thresher and Basking Shark.

Previous studies on plants^79,85^ and birds^78^ indicate that functionally and evolutionarily distinct species are disproportionately utilised by humans. Our work corroborates these earlier findings for birds and further shows that the same is true for sharks. Sets of bird and shark species selected based on functional irreplaceability scores highlight more species threatened by exploitation than selecting species at random. These findings support earlier work that suggests that functionally irreplaceable species are targeted by hunting^86,87^, fishing^86,87^ and the pet trade^88^. Beyond targeted fishing and exploitation, incidental catch is also a problem for many marine species^50,89^. Transitioning to the sustainable use and protection of species that are currently overexploited is essential to maintain functionally irreplaceable species and the benefits provided to people through ecosystem services.

Over the last two decades, the EDGE metric has galvanised conservation action by prioritising highly evolutionarily distinct and threatened species for practical conservation^7,20^. Our results build on this foundation by demonstrating how the new FIRE metric can effectively prioritise species to avert the greatest impending losses of functional diversity. The erosion and reorganisation of ecological assemblages is already widespread^90–92^, driven by rapid declines of species populations, particularly those with unique and vital ecological roles ^90–93^. Using a trait-based approach to prioritise species with distinct phenotypes and associated functional roles, the FIRE metric can help to bring a much-discussed but perennially overlooked dimension of biodiversity to the forefront of conservation planning, policy and action. Targeting this functional dimension is critical to maintain the complex and resilient ecosystems needed to support human wellbeing now and into the future^13^.

## 4. Methods

### Functionally Irreplaceable with Risk of Extinction (FIRE) calculations

The Functionally Irreplaceable with Risk of Extinction (FIRE) metric identifies species whose effective conservation will avert the greatest expected losses of functional diversity. To do this, for each species, FIRE combines two components: (i) functional irreplaceability, the expected unique contribution of a given species to overall trait space into the future, based on the extinction risk of all other species and (ii) the extinction risk of the species.

#### Species coverage of trait space

One way to measure functional irreplaceability, the property we are trying to maximise through our prioritisation scheme, is to quantify the proportion of TPD trait space uniquely covered by a species. As a first step it is necessary to find the hypervolume of trait space where a given TPD summed across all species is non-zero. More generally, however, we can calculate functional richness as;

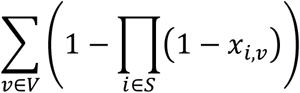

where 𝑥_𝑖,𝑣_ is the probability that species 𝑖 is present in a section 𝑣 of trait space, 𝑆 is the set of all species and 𝑉 is the set of all possible parts of trait space (these could be voxels in n-dimensional space). Our metric can be calculated as a binary or non-binary measure. In the binary measure we constrain 𝑥_𝑖,𝑣_ to be either 1 or 0. In the non-binary measure, we allow 𝑥_𝑖,𝑣_ to take any value between 0 and 1. These intermediate values may be based on uncertainty and/or intraspecific variability, which may in turn be connected to the total abundance of the species where such data are available. In cases where we assume some degree of intraspecific variation in species traits, each species occupies multiple voxels in trait space and the sum of occupancy across all voxels will be greater than 1. The equation can be adapted to incorporate extinction risk 𝐸_𝑖_ as a probability of extinction for each individual species 𝑖 ∈ 𝑆 to give an expression for the expected future hypervolume of trait space.

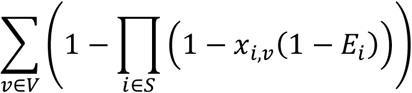

#### Irreplaceable species-specific trait space (functional irreplaceability)

To measure the irreplaceability of a species based on how much functional diversity it represents, we considered the extinction risk of all other species overlapping with the trait space of the focal species to reflect the expected unique occupancy of trait space in the future. If the focal species overlaps in trait space with few other species that are at high risk of extinction, there will be a moderate likelihood that this focal species will be the sole inhabitant of that region of trait space in the future. In contrast, if the focal species overlaps in trait space with many other species or with species that have low risk of extinction, then it is very unlikely that the focal species will be the sole inhabitant of that region of trait space in the future. The functional irreplaceability can be expressed mathematically as follows for species 𝑗 ∈ 𝑆

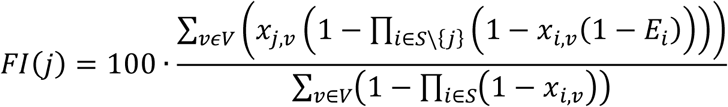

The numerator captures the amount of trait space that in the future, given extinction risk of all species other than the focal species, will rely on the survival of the focal species to remain covered. The denominator and factor of 100 expresses this as a percentage of total trait space. Because of the numerator in particular, this equation is computationally expensive to evaluate for all species. A naive implementation would have computational time complexity proportional to the size of the trait space multiplied by the square of the species richness. Furthermore, the whole calculation would still ideally need to be repeated to capture an uncertainty distribution propagating the error in all the Ei values. Our solution for calculating functional irreplaceability is to use the repeats required to capture uncertainty as part of a monte carlo process to simulate the result that the above equation captures in precise probabilistic terms. This requires three steps and is performed on every species (summarised in Fig. 1 of the main text).

Step 1: Calculate the volume of trait space occupied by all species in the analysis.

Step 2: Stochastically simulate the removal of species based on their extinction risk but with the constraint that the focal species does not go extinct (i.e., ignore the extinction risk of the focal species for this step). We followed the Gumbs et al.^20^ weighting of IUCN Red List categories^94^ to estimate probability of extinction for each species; IUCN Red List categories were converted to represent probability of extinction based on a continuous distribution of extinction probabilities, where the median of each Red List category was defined as, CR = 0.97; EN = 0.485; VU = 0.2425; NT = 0.12125; LC = 0.060625. Species that were not assessed or had IUCN Red List category listed as Data Deficient, were randomly sampled from the entire distribution of extinction probabilities. Using these probabilities, we ran simulations to determine whether species were removed from the pool (i.e., went extinct).

Step 3: Calculate a post-extinction trait space with the focal species present, and again with the focal species removed. Calculate the difference between these two trait space volumes, to give the species of interest’s distinct contribution to trait space. Divide the result by total trait space volume (step 1) and multiply this by 100 to get a percentage of the unique trait space occupied by the focal species resulting in a functional irreplaceability value.

#### Avertable loss (FIRE)

Step 4: To capture the avertable loss of irreplaceable functional diversity we multiplied the functional irreplaceability of each species (from step 3) by its probability of extinction to quantify the expected loss of functional diversity given the extinction risk of the species (i.e., the species’ FIRE value). We calculated risk of extinction by following the Gumbs et al.^20^ weighting of IUCN Red List categories^94^ to estimate the probability of extinction for each species. The full equation for FIRE of species 𝑗 ∈ 𝑆 is thus

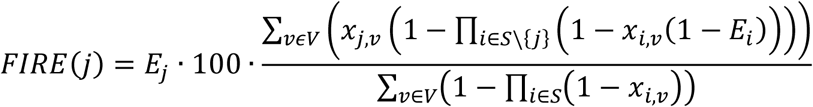

##### Final functional irreplaceability and FIRE values

We first perform step 1, then calculate functional irreplaceability and FIRE (perform steps 2-4 from above) over 1000 repeat iterations to capture a distribution of uncertainty in the extinction scenarios. The final functional irreplaceability and FIRE values are given by the median value of the sampled distribution.

### Case study 1: FIRE for the world’s birds

#### Data collection

We collated published data for extinction risk, traits and evolutionary history of birds (Aves). To taxonomically match species information from different datasets, we used nomenclature from the latest Birdlife checklist^95^ (accessed 10/2024), which included 11005 species. All database manipulation and analyses were performed using R statistical software (v. 4.3.1)^96^.

##### Extinction risk data

We use the IUCN Red List of Threatened Species (v. 2025.1)^2^ to obtain information of species extinction risk. This includes assessment data for 10975 species, including 10936 data sufficient species (Least Concern, LC; Near Threatened, NT; Vulnerable, VU; Endangered, EN; or Critically Endangered, CR) and 39 categorised as Data Deficient (DD). The remaining 30 species in our dataset are Not Evaluated (NE) on the Red List. To calculate a species probability of extinction we generate a continuous distribution of extinction probabilities following methods outlined by Gumbs et al.^20^, where the median of each Red List category was defined as, CR = 0.97; EN = 0.485; VU = 0.2425; NT = 0.12125; LC = 0.060625. Species probability of extinction was randomly selected from the distribution of extinction probability values within their Red List categorisation. Where species were categorised as DD or NE their extinction probability was randomly selected from the whole distribution of extinction risk values.

##### Evolutionary history data

We used the EDGE2 protocol^20^ to calculate evolutionary distinctiveness (ED; the expected contribution of a species to overall evolutionary history given the extinction risk to all other species) and Evolutionarily Distinct and Globally Endangered (EDGE; the amount of expected loss of evolutionary history that can be conserved by averting the extinction of a given species) scores for all birds. Evolutionary distinctiveness and EDGE are the phylogenetic equivalents of functional irreplaceability and FIRE, and we used the same weightings of extinction risk for EDGE and FIRE calculations. ED and EDGE scores were calculated as the median scores from a sample of 1000 phylogenetic trees generated from Jetz et al.^97^ phylogeny, where species absent from the phylogenies were inserted based on taxonomy using the same approach as Gumbs et al.^41^.

##### Trait data

As trait data for birds, we used eight continuous morphological measurements that have been demonstrated to reliably indicate the functional roles and trophic statuses of birds on a global scale^37^: (1) total beak length (from the tip to the skull), (2) beak length to the nares, (3) beak width and (4) depth (at the nares), (5) wing length, (6) secondary length (length from the carpal joint to the first of the wings secondary feathers), (7) tail length and (8) tarsus length. Data for all eight traits were sourced from the AVONET database^33^. We also sourced information on species body mass from AVONET for use in a sensitivity test. We investigated the relationship between traits and extinction risk using Spearman’s rank correlation.

#### Quantifying trait space

##### Principle component analysis

The eight morphological traits were all log-transformed and then centred and scaled to a mean of zero and unit variance. A Principle Components Analysis (PCA) was undertaken using the eight log-transformed and scaled traits and the first three PCA axes taken to build the trait space. We used three axes as they explained over 90% of the total variance and resulted in a more manageable number of species (compared to when using four PCA axes) with the maximum functional irreplaceability value. The latter is because, as trait space dimensionality increases, there is more trait space available in which to measure differentiation and thus in most cases, and keeping all other parameters equal, species become more functionally distinct from each other.

##### Multidimensional trait space

We use TPDs to account for intraspecific trait variability when describing functional space^34^. TPDs represent each species in functional trait space as a multivariate kernel density estimation. We approximated TPD values using the measured traits available in the analysis and an estimated variability using the TPD package (v. 1.1.0)^34^. Each species occupancy is estimated in the same field defined by the three axes of the PCA, with each grid divided into 20 equal parts. Trait space is held constant following removal of species in subsequent analyses.

##### Functional irreplaceability/FIRE

We apply the methods stated above to calculate functional irreplaceability and FIRE values for all birds using the non-binary method to calculate species occupancy of trait space (where the species proportional probabilistic occupancy of each voxel is calculated). Following the EDGE2 protocol^20^ and applying it to our FIRE metric, we defined sets of priority FIRE birds as being those that are both assessed as threatened on the IUCN Red List and above the median, 75th percentile, and 95th percentile of functional irreplaceability with 95% confidence (i.e. in 95% or more of the iterations of FIRE calculation; Data S1 and S2).

##### Accounting for outliers in trait space

The five kiwi species (Apterygiformes) represent extreme outliers in the global bird morphological trait space^37^ and can have substantial impact on functional diversity analyses if not accounted for^61^. Here, to mitigate this effect we first built the trait space and calculated functional irreplaceability for all species with the kiwis included, storing the functional irreplaceability values for the five kiwi species. We then removed the five kiwi species and re-built the trait space (first re-scaling the traits and re-running the PCA). This second trait space was used to calculate functional irreplaceability for all other bird species (i.e., all species other than the kiwis). The functional irreplaceability values for the kiwis were then added to these values prior to FIRE values being calculated for all birds. This process means that, when the second trait space is analysed, the kiwis are not included in the species removed based on their extinction risk (i.e., Step 2 above).

However, this is unlikely to be an issue given they occupy an entirely distinct area of trait space and thus will not influence the functional irreplaceability of any other species.

##### Trait space extinction projections

To understand the future impact of species extinctions on global trait space, we estimated the proportional loss of each voxel in trait space. Future occupancy of voxels in trait space is calculated as a function of current occupancy of species in trait space and extinction probabilities. To estimate this, we calculate the probability that at least one species remains in a given voxel, after accounting for extinction probabilities, and divide this by the probability that at least one species currently occupies that voxel:

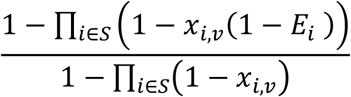

Where 𝑥_𝑖,𝑣_ is the probability that species 𝑖 is present in voxel 𝑣 of trait space, 𝐸_𝑖_ is the probability of extinction for species 𝑖 and 𝑆 is the set of all species. Each future projection is calculated 1000 times to reflect the distribution of species extinction predictions. These predictions were mapped onto the three principal component axes of trait space to visualise the impact of extinction on different areas of trait space and identify certain trait combinations that may be vulnerable to extinction.

### Avian functional irreplaceability

We used functional irreplaceability scores to identify bird species with little or no functional redundancy, including those with no overlap with other species in our trait space. To explore the relationship between functional irreplaceability and extinction risk, we used Welch’s t-test to compare the mean functional irreplaceability of threatened (i.e. VU, EN, CR on the Red List) and non-threatened (LC and NT on the Red List) species. We selected the species in the top 5% highest functional irreplaceability scores (following Safi et al. 2013^98^), and calculated the proportion that are threatened, and compared this qualitatively with the overall proportion of threatened birds, derived from the 2025.1 version of the IUCN Red List^2^.

### Unique and convergent FIRE priorities

To compare convergent and unique conservation priorities between FIRE and the phylogenetically-informed EDGE approach, we matched the bird species in our FIRE dataset with those with available EDGE data. To achieve this, we used existing EDGE2 scores for birds^41,99^, updated for the 2025.1 Red List extinction risk data^100^. We limited our comparisons of FIRE and EDGE data to species for which both data were available (N = 10971). To quantify the relationship between avian functional irreplaceability and its phylogenetic equivalent, ED, we used Spearman’s rank correlation. We also used Welch’s t-test to test whether species in monotypic families had higher functional irreplaceability than birds in general. We again followed Safi et al.^98^ in highlighting species with top 5% highest FIRE scores as top priorities, and restricted this set to species in a threatened or Extinct in the Wild Red List category. Using this set of priority species, we identified the number of species also in the top 5% threatened species based on EDGE scores. We also calculated what proportion of these priority species qualify as priority EDGE species (i.e. above median EDGE with 95% confidence and threatened with extinction^20^), to identify species not currently captured by a primary phylogenetically-informed prioritisation. To explore how well the priority FIRE species are represented by other conservation mechanisms, we calculated how many species trigger Key Biodiversity Areas^101^ (KBAs), and used published estimates of range overlap with KBAs^102^ to identify species potentially underrepresented in the global KBA network. Lansley et al.^102^ suggested that species with <8% of their area of habitat^103^ (AOH; IUCN range maps trimmed to suitable habitat and elevation) covered by KBAs may not be effectively represented in priority areas for maintaining biodiversity.

### Case study 2: Calculating FIRE for sharks

#### Data collection

We collated published data for extinction risk, traits and evolutionary history of sharks (Selachimorpha). We limited our study to sharks, omitting other clades of Chondrichthyes (Batoidea and Holocephali), due to limited availability of comparable data for several key traits, including body shape. To taxonomically match species information from different datasets, we used nomenclature from FishBase^64^ (accessed 11/2023), which included 557 species.

##### Extinction risk data

We use the IUCN Red List of Threatened Species (v. 2023.1)^104^ to obtain information of species extinction risk. This includes assessment data for 540 species, including 467 data sufficient species (Least Concern, LC; Near Threatened, NT; Vulnerable, VU; Endangered, EN; or Critically Endangered, CR) and 73 categorised as Data Deficient (DD). The remaining 17 species in our dataset are Not Evaluated (NE) on the Red List. To calculate a species probability of extinction we generate a continuous distribution of extinction probabilities following methods outlined by Gumbs et al.^20^, where the median of each Red List category was defined as, CR = 0.97; EN = 0.485; VU = 0.2425; NT = 0.12125; LC = 0.060625. Species probability of extinction was randomly selected from the distribution of extinction probability values within their Red List categorisation. Where species were categorised as DD or NE their extinction probability was randomly selected from the whole distribution of extinction risk values.

##### Evolutionary history data

We used the EDGE2 protocol^20^ to calculate evolutionary distinctiveness and EDGE scores for all sharks. ED and EDGE scores were calculated as the median scores from a sample of 1000 phylogenetic trees generated from the Stein et al.^105^ tree distribution of 10,000 Chondrichthyes trees, where species absent from the phylogenies were inserted based on taxonomy using the same approach as Gumbs et al.^41^.

##### Trait data

We collated eight traits for our analysis (Data S4), obtained from FishBase^64^ and Sharks of the World: A Complete Guide^106^. We selected traits based on data availability and representation of functional guilds^107^ (Table S1). Our traits were selected to capture a broad suite of traits that are important to describing species’ roles within an ecosystem reflecting life history strategies, habitat use, trophic interactions and morphological variation (Table S1)^108,109^. To measure body shape, we modify methods shown in Siders et al.^108^, which uses Elliptical Fourier analysis (EFA) to represent variation in shark body shape. We collected 515 illustrated images from Sharks of the World: A Complete Guide to measure the variation in shark body shape using EFAs. These images were first converted into outlined silhouettes using photoshop (v. 23.5.5)^110^. Using the R package Momocs (v. 1.4.1)^111^ each outline was scaled and centred on a matrix (Fig. S5) and EFA was performed on the outlines, using 16 harmonics (Fig. S6). Results of the EFA were condensed using Principal Component Analysis (PCA; Data S5). PC1 and 2 collectively explain 62% of total variation and were used to represent body shape for our analysis (Fig. S7), PC1 represents 43% variation and PC2 represents 19% variation.

#### Imputing missing data

Our compiled database contained 557 species. To complete missing values in the trait database, we imputed missing data using the missForest function in the missForest R package (v. 1.5)^51^. This method uses a random forest trained on the observed values of the trait matrix to impute missing values. It can be used to predict both continuous and categorical data.

To test the performance of imputation we limited the dataset to only include species with complete trait data and simulated incomplete trait data by artificially removing trait data. We randomly removed data from each trait, with the percentage of the trait removal equal to the percentage of the trait data missing from our full database. Simulated removal of data was repeated 100 times to get a variable combination of missing values. We then imputed the missing data using missForest and calculated each species functional irreplaceability 100 times, and calculated this 100 times from the unimputed dataset. We compared the median functional irreplaceability of each species from the complete dataset and the simulated imputed dataset using Spearman’s rank correlation to test the robustness of imputation using missForest for the observed proportions of missing data.

To examine the robustness of FIRE rankings to the inclusion of species with missing data, we calculated FIRE rankings for the set of species with complete trait data and compared these to the rankings of the species when FIRE was calculated for all species (including imputed data). To examine the robustness of priority species identified, we calculated the number and proportion of species that were in the top 10% and 20% of both rankings. We then used Spearman’s rank correlation to examine the relationship between the absolute change in rankings for each species between the two datasets, and the priority ranking of each species in the FIRE rankings restricted to species with complete trait data. If there is no relationship, this indicates that high priority rankings are similarly unstable as low-priority rankings, whereas a positive correlation would indicate greater stability in the ranks of high priority species.

To assess relationships between species traits and extinction risk, we used Spearman’s rank correlation to quantify relationships with extinction risk and continuous trait (body length, trophic level, maximum depth, growth ratio, body shape variation PC1 and body shape variation PC2). For categorical traits (habitat preference and reproductive guild), we used Kruskal-Wallis tests to test for differences in extinction score among trait categories.

#### Quantifying trait space

##### Distance matrix

Prior to quantifying a distance matrix, we log transformed, scaled and centred continuous traits to ensure normal distribution and prevent overweighting. We calculated a dissimilarity distance matrix (Gower’s distance) using the function gawdis (v. 0.1.5)^14^. This approach prevents disproportionate weighting of categorical traits to produce multi-trait dissimilarity with more uniform contributions of different traits^14^.

##### Principal coordinate analyses

Using the dudi.pco function in the ade4 package (v. 1.7.22)^112^ we summarised the functional dissimilarity matrix with PCoA to quantify distances in trait space between species. Multivariate trait space was built by extracting the first four PCoA axes. To assess how imputation was distributed across trait space, we calculated the proportion of imputed data for each species and summarised imputation density across trait space. We calculated the average proportion of imputed data along the first two PCoA axes by summarising the mean of each point along both PCoA axes.

##### Multidimensional trait space

We approximated TPD values using the measured traits available in the analysis and an estimated variability using the TPD package (v. 1.1.0)^34^. Each species occupancy is estimated in the same field defined by the four axes of the PCoA, with each grid divided into 25 equal parts. Trait space is held constant following removal of species in subsequent analyses.

##### Functional irreplaceability/FIRE

We apply the methods stated above to calculate functional irreplaceability and FIRE values for all sharks using the binary method to calculate species occupancy of trait space (either a species occupies a voxel (1) or it does not (0)). Following the EDGE2 protocol^20^ and applying it to our FIRE metric, we defined sets of priority FIRE sharks as being those that are both assessed as threatened on the IUCN Red List and above the median, 75th percentile, and 95th percentile of functional irreplaceability with 95% confidence (i.e. in 95% or more of the iterations of FIRE calculation; Data S2).

### Relationship between FIRE and EDGE

We used Pearson’s correlations to examine the relationship between functional irreplaceability and evolutionary distinctiveness scores and between FIRE and EDGE scores. To determine species that are of particularly high priority for conserving irreplaceable functional and phylogenetic diversity we identified species that are in the top 5% of both functional irreplaceability and evolutionary distinctiveness as well as FIRE and EDGE.

To better understand patterns of high functional irreplaceability in highly speciose groups (i.e. those with many close evolutionary relatives) we investigated the distribution of three species— the Gummy Shark (*Mustelus antarcticus*), Rig (*Mustelus lenticulatus*) and Southern Sleeper Shark (*Somniosus antarcticus*)—and their congeners in trait space. We plot each dimension of trait space and highlight the three species and their congeners in trait space (Fig. S4).

### Functional irreplaceability and exploitation

We identified exploited bird and shark species using the IUCN Red List threats classification scheme (https://www.iucnredlist.org/resources/threat-classification-scheme)^2^, where threat type was listed as Biological resource use. We consider a species intentionally exploited when the species being assessed is the target, as codified under Intentional use, Intentional use: subsistence/small scale and Intentional use: large scale. Unintentionally exploited species were those indicated as being impacted by Unintentional effects, Unintentional effects: subsistence/small scale and Unintentional effects: large scale. Species that were listed as both intentional use and unintentional effects were not included in the unintentionally exploited categorisation.

To explore the potential of different prioritisation strategies at safeguarding intentionally exploited species, we selected 10%, 20%, 30%, 40% and 50% of all bird and shark species, weighted by different prioritisation measures: evolutionary distinctiveness score; functional irreplaceability score and randomly selected species. We repeated this 1000 times for each approach and compared the number of intentionally exploited species captured by each approach for each sample size. To test if prioritisation strategies differed significantly at safeguarding intentionally exploited species we used ANOVA with Tukey’s Honestly Significant Difference post hoc test to test the pairwise difference between strategies.

To consider species that are unintentionally exploited we compared functional irreplaceability and evolutionary distinctiveness scores of species using t-tests. To investigate the potential impact of human exploitation on shark functional diversity and phylogenetic diversity, we removed species categorised as i) intentionally exploited and ii) both intentionally and unintentionally exploited to identify the loss of trait space and cumulative years of evolutionary history. We compared the loss of trait space and evolutionary history in exploited species to a randomly selected sample of species of the same number.

## Supplementary material

### Supplementary text

#### S1 Text | Bird functional irreplaceability; hummingbirds as a case study

Inspecting the list of the most functionally irreplaceable birds indicates that there are numerous hummingbirds included: 12 of the top 50 functionally irreplaceable birds are hummingbirds. Hummingbirds as a group are known to be particularly functionally distinctive^1,2^. However, within hummingbirds, while some of these species are widely recognised to be functionally unique species (e.g., sword-billed hummingbird *Ensifera ensifera* – the only species with a beak longer than its body, excluding the tail), the inclusion of some of the other species in the list of most functionally irreplaceable birds is arguably less intuitive. To check that our analytical approach was working as intended, we inspected the location of the most irreplaceable hummingbird species in the 3-dimensional global bird trait space (the trait space excluding the five kiwi species; Fig. A). In Figure A hummingbirds (Trochilidae) are shown as red points with the six most functionally irreplaceable hummingbird species shown as blue points, and all other bird species as black points. Two things are apparent when inspecting Fig. A: First, as expected, hummingbirds as a group are relatively distinct relative to all other birds, at least in this 3-dimensional trait space. Second, the hummingbirds assessed to be the most functionally irreplaceable in our analyses are clearly located on the edge of trait space, thus indicating that our analyses are working as expected. It is likely that, while the individual traits of some species may not make some species appear particularly functionally distinct to the naked eye (i.e., they may not have extreme individual trait values such as a very long bill), the use of PCA better highlights species that have rare combinations of trait values (i.e., species that are relatively distinct in PCA space). A similar process is observed with certain palm swift species, where swifts in general are located on the edge of trait space, and then certain palm swift species – those that have rare combinations of trait values – are located on the edge of this group and thus occupying the extremities of trait space. As a final sense check, we highlighted the location of the species with the top 100 functional irreplaceability values in the 3-dimensional global bird trait space (Fig. B); this plot clearly shows that the species are, as expected, located towards the edges of trait space.

**Fig. A.**
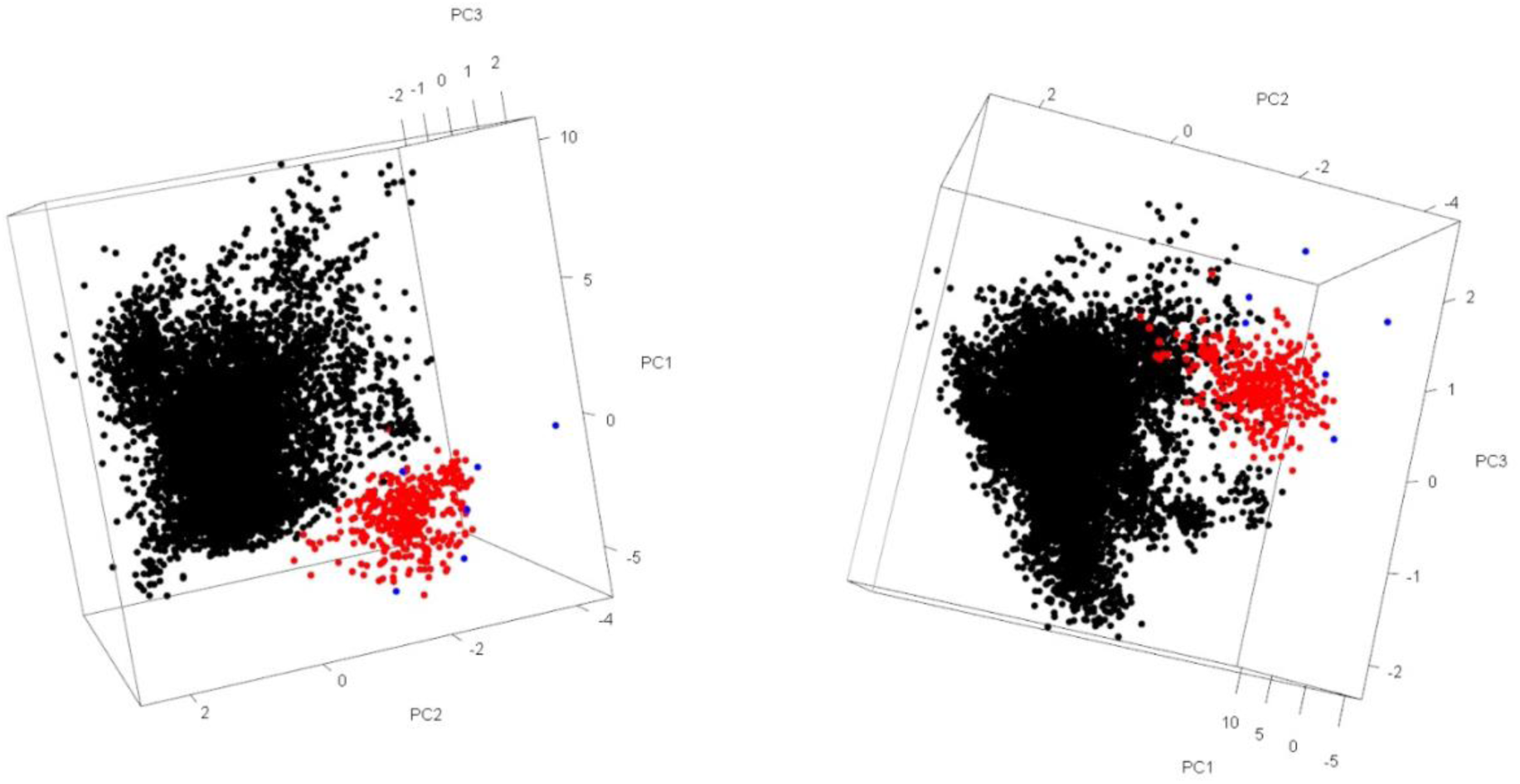
| The location of hummingbirds (red points; n = 360) in the global bird trait space. The six hummingbird species with the highest functional irreplaceability scores (*Coeligena orina*, *Discosura letitiae*, *Phaethornis yaruqui*, *Pterophanes cyanopterus*, *Ensifera ensifera*, *Chaetocercus berlepschi*) are shown as blue points and all other species (n = 10,634) as black points. The trait space is that used in the main analyses, built after excluding the five kiwi species. Both plots show the same trait space at different rotations.

**Fig. B.**
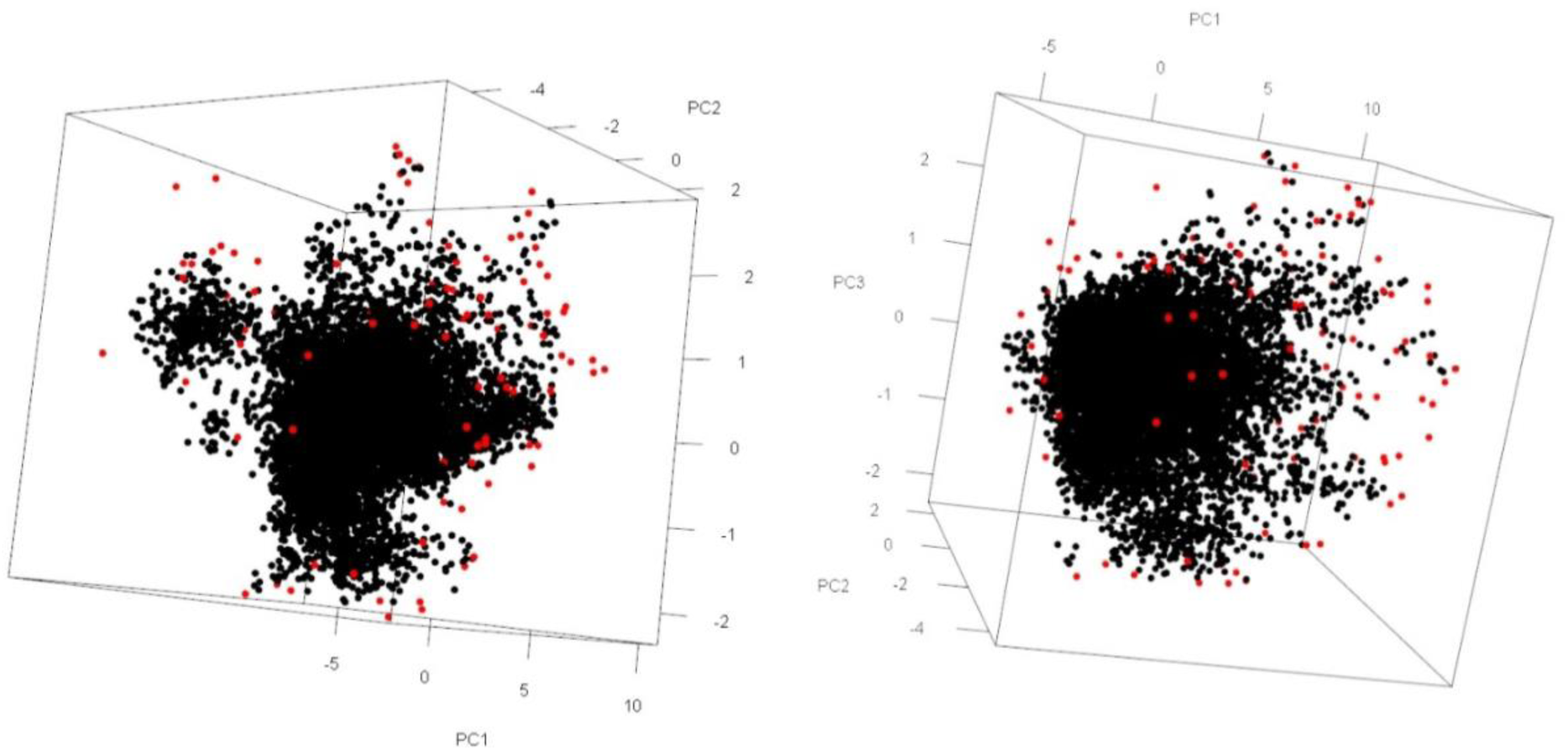
| The location of the 100 most functionally irreplaceable birds (red points) in the global bird trait space. All other species (n = 10,900) are shown as black points. The trait space is that used in the main analyses, built after excluding the five kiwi species. Both plots show the same trait space at different rotations.

#### S2 Text | Shark functional irreplaceability and extinction risk

##### Current unique irreplaceability and FIRE values

Incorporating functional complementarity [i.e. considering the extinction risk of species closely associated in trait space] into the functional irreplaceability calculations will change the consequences of each species’ functional irreplaceability score, depending on their relation in trait space with other species, and the extinction probability of those species. To test the correlation between current unique irreplaceability and our measure of functional irreplaceability based on expected loss, we also calculate current irreplaceability as the area of current trait space uniquely occupied by a species. We calculated the proportion of each species’ functional irreplaceability score that was due to their unique contribution to current trait space compared with the proportion gained through extinction scenarios (Fig. A).

**Fig. A.**
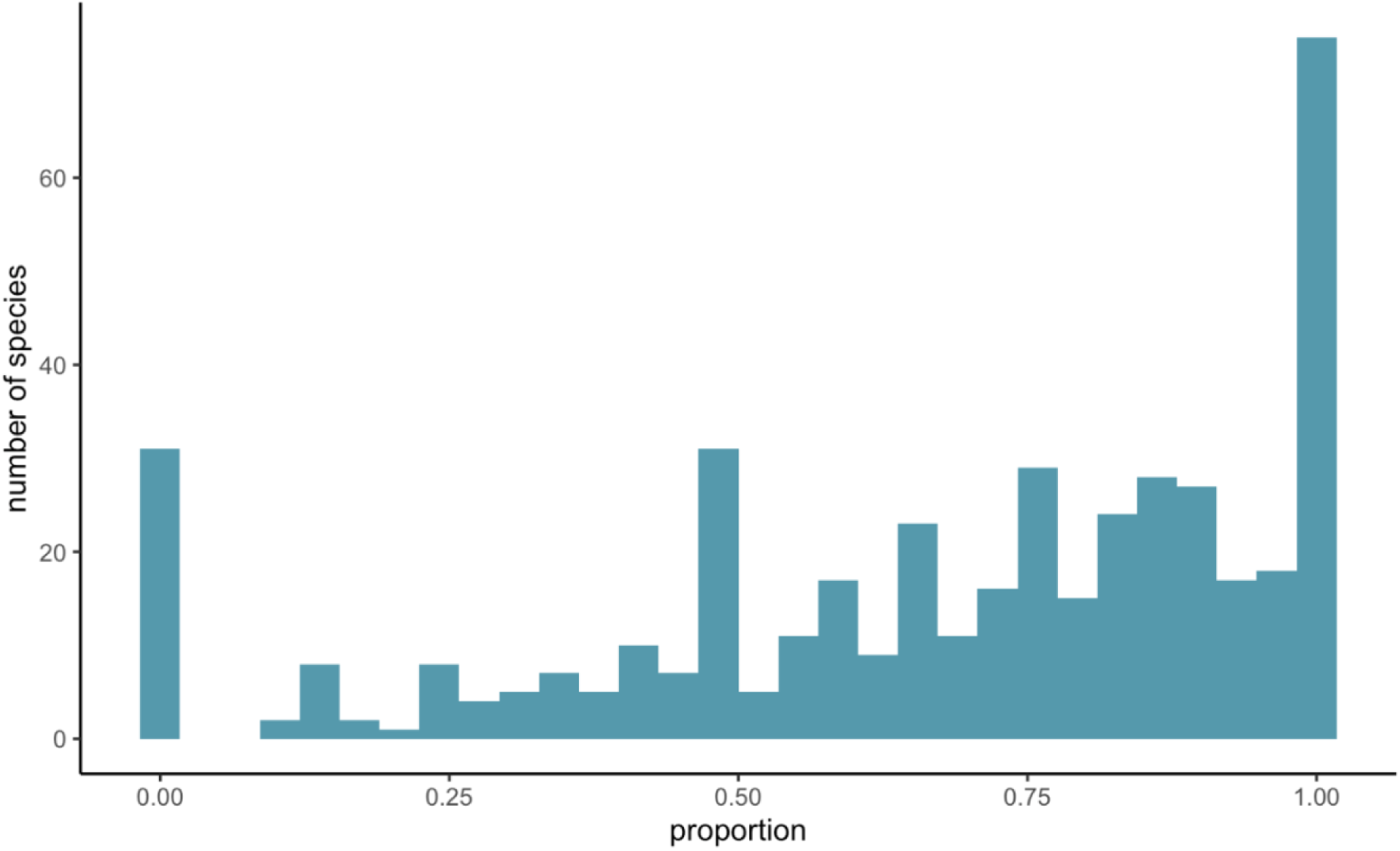
| Proportion of unique functional distinctiveness of each species represented by current trait space occupancy. The proportion of each species functional distinctiveness score due to current trait space occupancy compared to future trait space occupancy. The median amount of functional distinctiveness represented by current trait space is 75% of the species functional distinctiveness score.

To ensure there is no overweighting of extinction probabilities, we calculated FIRE scores from these current unique irreplaceability scores rather than those from an extinction-based approach, and correlated those with the original FIRE scores. There was a strong correlation between functional irreplaceability and a measure of unique current contributions to trait space for each species (ρ = 0.63, p < 0.001; Fig. B).

**Fig. B.**
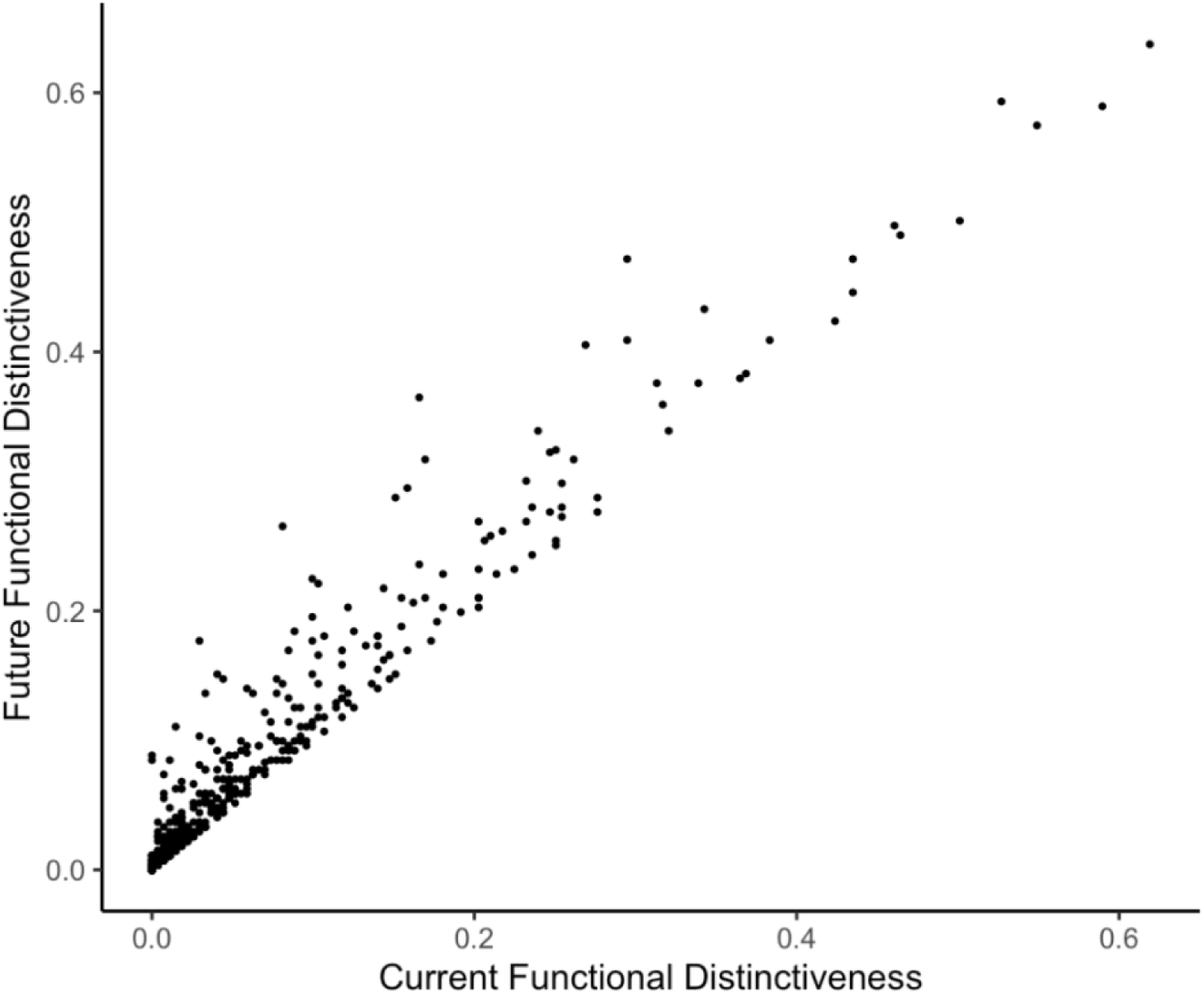
| Correlation between future and current functional distinctiveness. A scatterplot to compare the relationship between future functional distinctiveness (where functional complementarity is accounted for) and current functional distinctiveness (where all species are included in trait space when calculating functional distinctiveness). Correlations between future and current functional distinctiveness were significantly positive (Spearman’s correlation: r = 0.96, p < 0.001).

##### Comparing functional metrics

To investigate the variations between different metrics, we use Pearson’s correlation to compare functional irreplaceability against other approaches to quantifying species-specific contributions to overall functional diversity. We investigate correlations between functional irreplaceability, functional uniqueness (FUn), functional distinctiveness (FDis) and functional uniqueness measured in probabilistic trait space (FU). FU is calculated as the functional distance to the nearest neighbour of the species of interest, using the uniqueness function in the funrar package (v. 1.5.0)^3^. FDis is calculated as the average functional distance from a species to all the others in the given community, using the distinctiveness function in the funrar package (v. 1.5.0)^3^. FU is calculated as the mean overlap of species in probabilistic trait space, using the uniqueness function in the TPD package (v. 1.1.0)^4^. We compared our functional irreplaceability measure against other approaches to quantifying species-specific contributions to overall functional diversity, and found significant but only moderate positive relationships between them (Fig. C).

**Fig. C.**
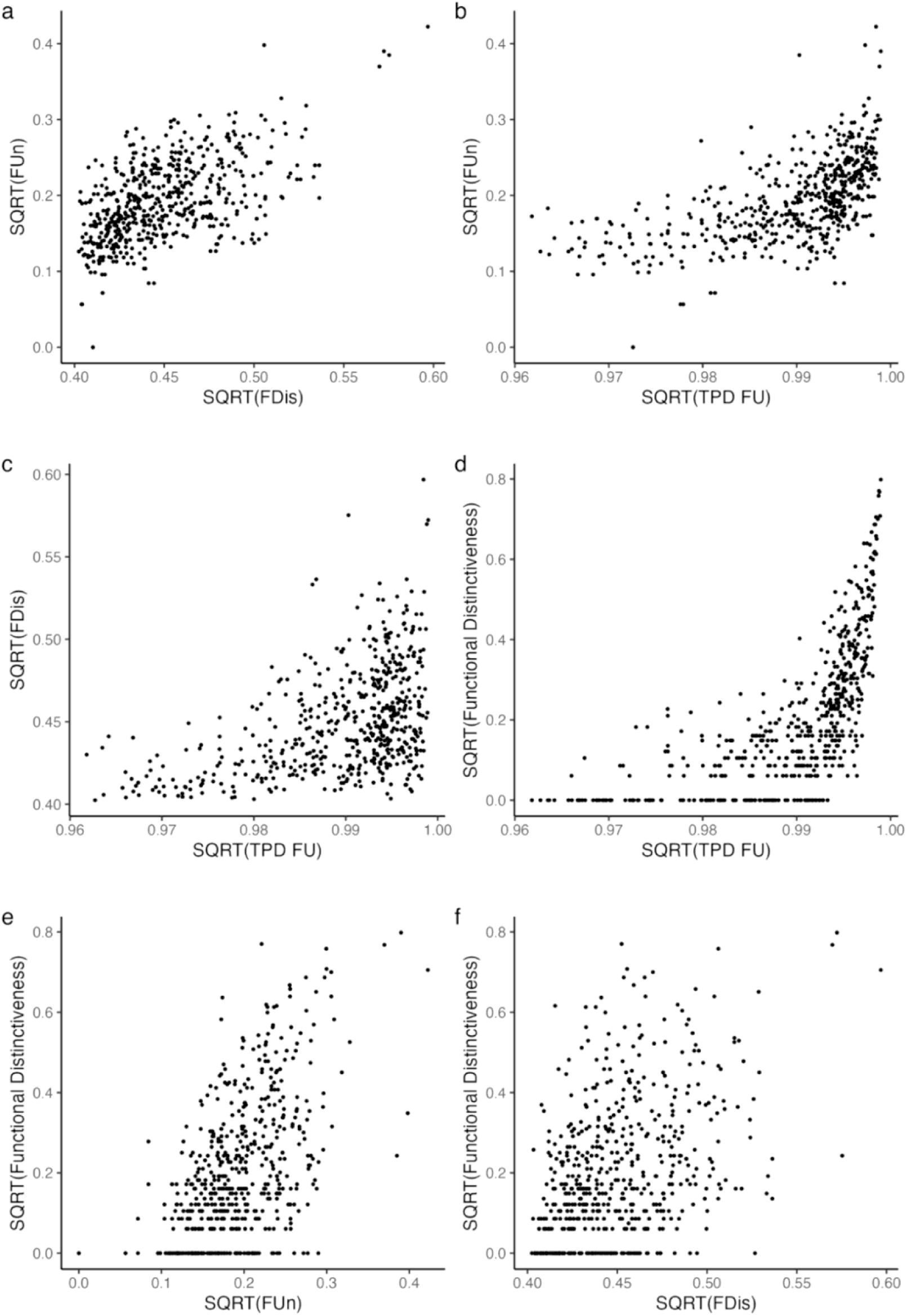
| Correlation between functional indices. Scatterplots to compare species-specific functional diversity indices between; a) Functional Uniqueness (FUn) and Functional Distinctiveness (FDis; Pearson’s correlation: r = 0.56, p < 0.001), b) Functional Uniqueness and Trait Probability Density Functional Uniqueness (TPD FU; Pearson’s correlation: r = 0.57, p < 0.001), c) FDis and TPD FU (Pearson’s correlation: r = 0.47, p < 0.001), d) TPD functional distinctiveness and TPD FU (Pearson’s correlation: r = 0.66, p < 0.001), e) TPD functional distinctiveness and FUn (Pearson’s correlation: r = 0.55, p < 0.001) and f) TPD functional distinctiveness and FDis (Pearson’s correlation: r = 0.43, p < 0.001).

##### Global shark functional irreplaceability

The top 5% of functionally irreplaceable shark species (28 spp.) together occupy 10.85% of unique trait space. Overall, 46% of the top 5% functionally irreplaceable species are threatened with extinction, compared with 30% of all sharks^5^. To estimate the projected impact of extinction on shark trait space, we calculated the proportion of trait space that would be lost with the extinction of all 167 currently threatened species (Vulnerable [VU], Endangered [EN], Critically Endangered [CR] on the IUCN Red List). We found that 17.18% (compared to an average random sample of 11.53%; sd = 1.24) of global shark trait space is at risk of extinction, potentially increasing to 23.58% (compared to an average random sample of 20.73%; sd = 1.58) if we were to also consider all Data Deficient [DD] and Not Evaluated [NE] species as at risk of extinction. The orders Hexanchiformes, Echinorhiniformes and Lamniformes have the highest average functional irreplaceability scores of 0.20% (sd = 0.021), 0.18% (sd = 0.001) and 0.18% (sd = 0.018) respectively (Fig. D). The orders with the lowest average functional irreplaceability score are Squatiniformes and Carcharhiniformes, which both have an average functional irreplaceability score of 0.06% (sd = 0.007 and 0.009, respectively) (Fig. D).

**Fig. D.**
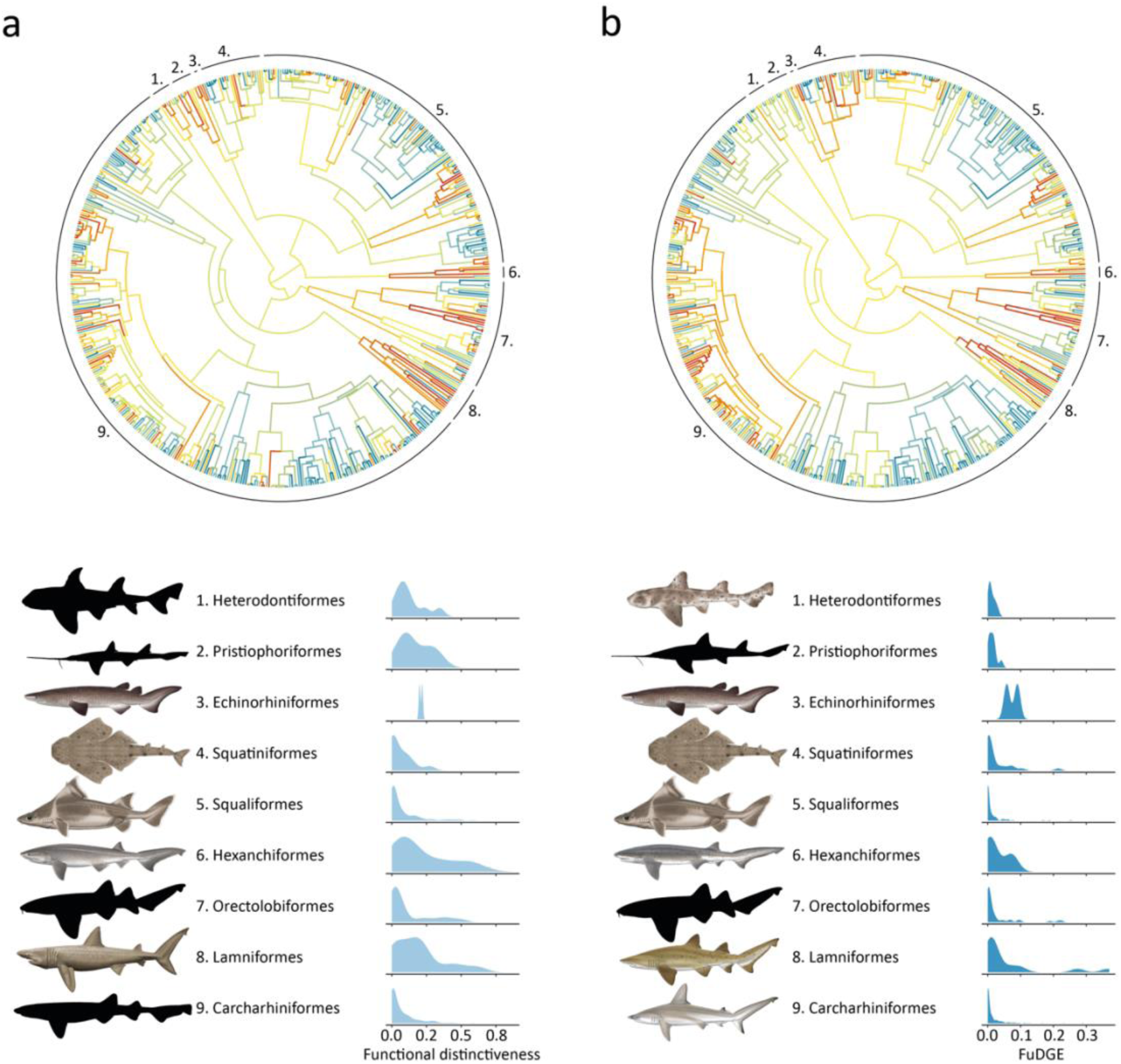
| Distribution of functional irreplaceability and FIRE across the tree of life. A phylogenetic tree of all sharks with branches coloured to represent the median (a) functional irreplaceability and (b) FIRE of all descendant species across the shark tree of life. Numbers indicate shark orders. Density plots represent the distribution of (a) functional irreplaceability and (b) FIRE scores in each order, both on log scale. Illustrations represent the shark species with the highest (a) functional irreplaceability and (b) FIRE score in their respective order. Shark species: (a) 1. Barred Bullhead Shark (*Heterodontus zebra*), 2. Warren’s Sixgill Sawshark (*Pliotrema warreni*), 3. Bramble Shark (*Echinorhinus brucus*), 4. Smoothback Angelshark (*Squatina oculata*), 5. Angular Roughshark (*Oxynotus centrina*), 6. Bluntnose Sixgill Shark (*Hexanchus griseus*), 7. Pacific Nurse Shark (*Ginglymostoma unami*), 8. Basking Shark (*Cetorhinus maximus*), 9. Japanese Swellshark (*Cephaloscyllium umbratile*), (b) 1. Horn Shark (*Heterodontus francisci*), 2. Anna’s Sixgill Sawshark (*Pliotrema annae*), 3. Bramble Shark (*Echinorhinus brucus*), 4. Smoothback Angelshark (*Squatina oculata*), 5. Angular Roughshark (*Oxynotus centrina*), 6. Broadnose Sevengill Shark (*Notorynchus cepedianus),* 7. Pacific Nurse Shark (*Ginglymostoma unami*), 8. Sand Tiger Shark (*Carcharias taurus*), 9. Scalloped Hammerhead (*Sphyrna lewini*). The phylogenetic tree was selected at random from the set of 1,000 phylogenetic trees used in this study to calculate EDGE scores. Illustrations: Marc Dando.

## Supplementary figures and tables

**Fig. S1.**
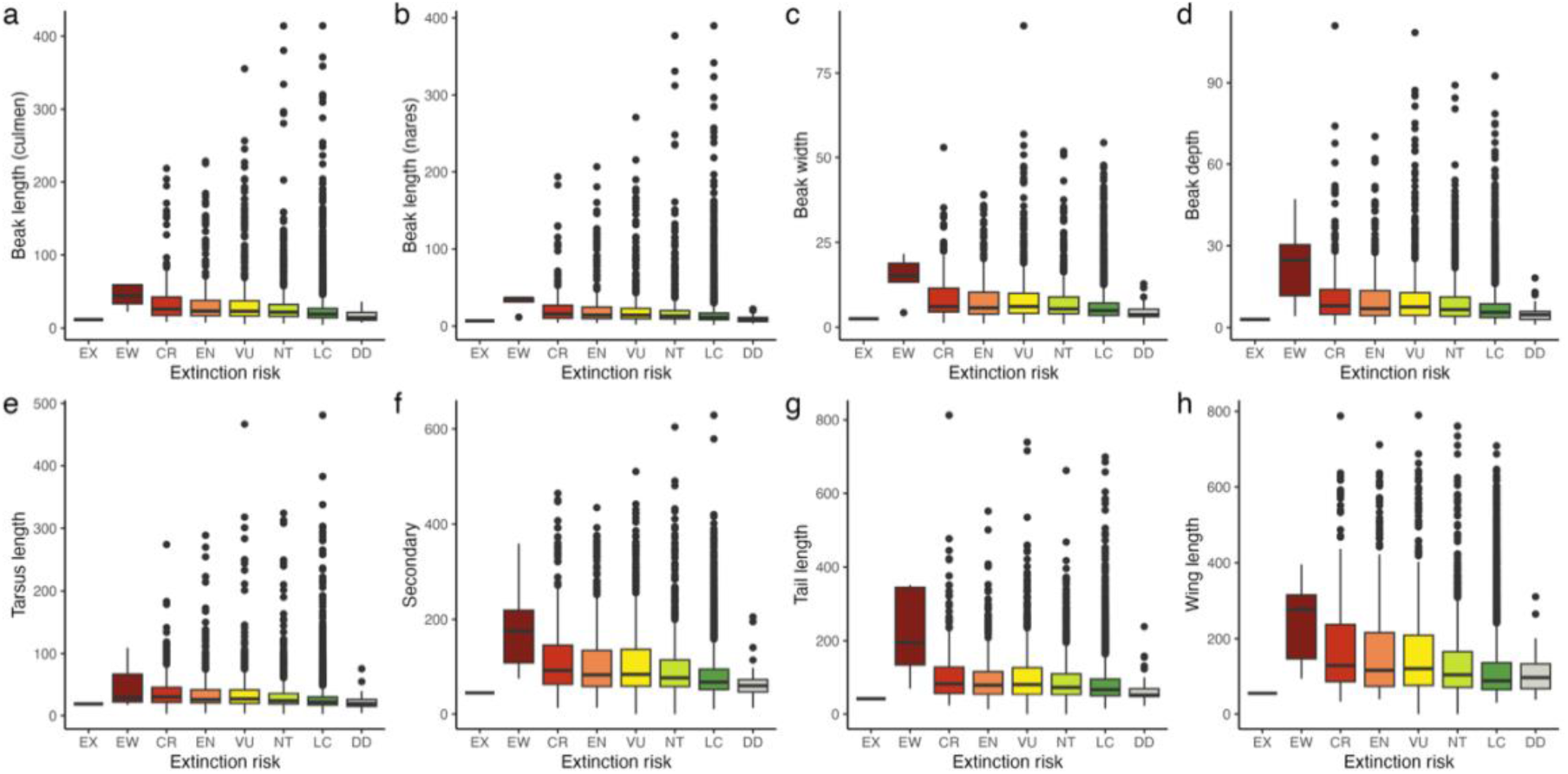
| Relationships between bird morphological traits and extinction risk. Variation in morphological trait values across IUCN Red List categories (EX = Extinct, EW = Extinct in the Wild, CR = Critically Endangered, EN = Endangered, VU = Vulnerable, NT = Near Threatened, LC = Least Concern, DD = Data Deficient). Traits include: (a) beak length (culmen), (b) beak length (nares), (c) beak width, (d) beak depth, (e) tarsus length, (f) secondary length, (g) tail length, and (h) wing length. Boxplots represent the range, interquartile range and median with points denoting outliers.

**Fig. S2.**
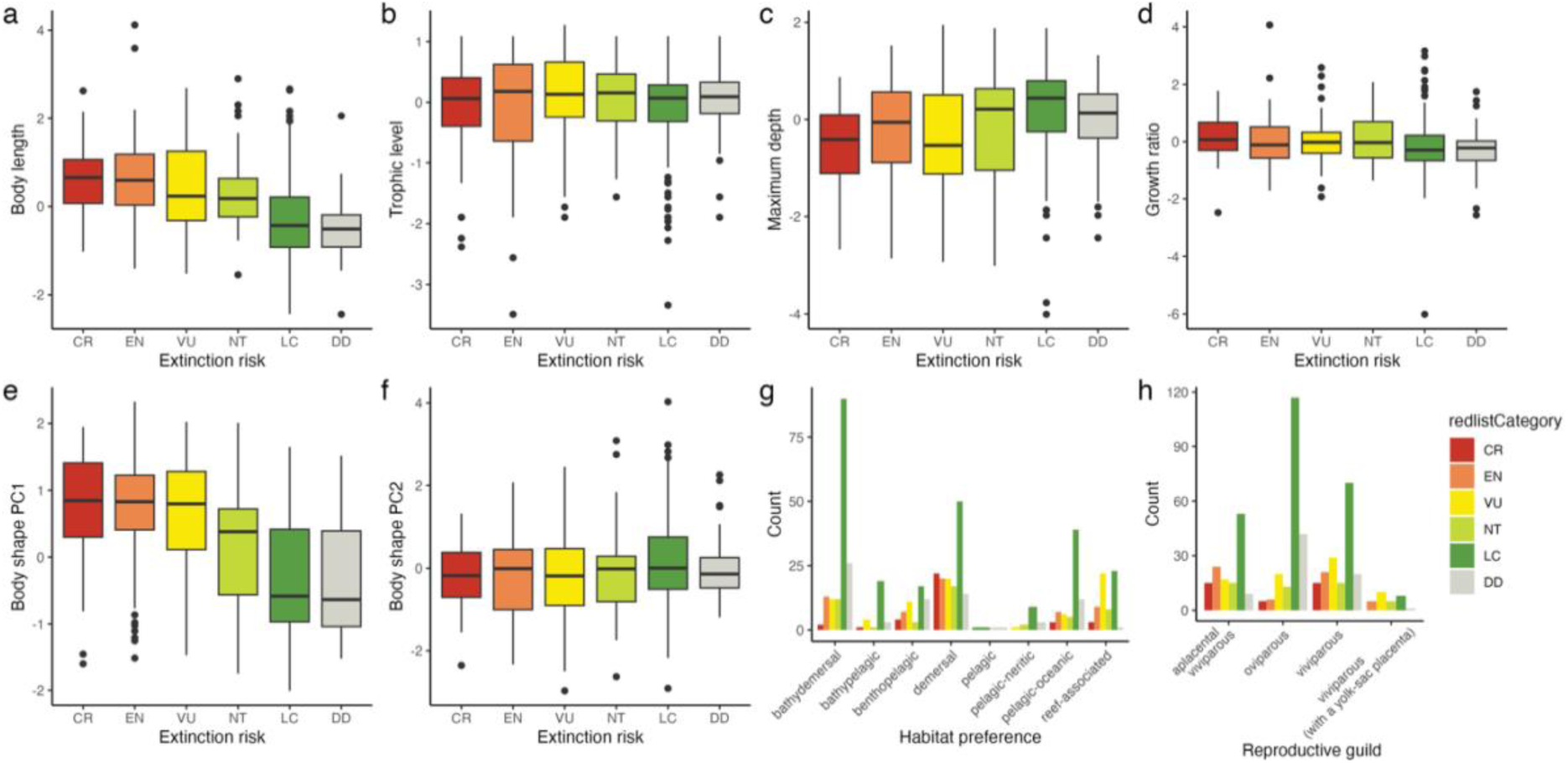
| Relationships between shark traits and extinction risk. Variation in trait values across IUCN Red List categories (CR = Critically Endangered, EN = Endangered, VU = Vulnerable, NT = Near Threatened, LC = Least Concern, DD = Data Deficient). Traits include: (a) body length, (b) trophic level, (c)maximum depth, (d) growth ratio, (e) body shape variation PC1, (f) body shape variation PC2, (g) habitat preference, and (h) reproductive guild. Boxplots (a-f) represent the range, interquartile range and median with points denoting outliers.

**Fig. S3.**
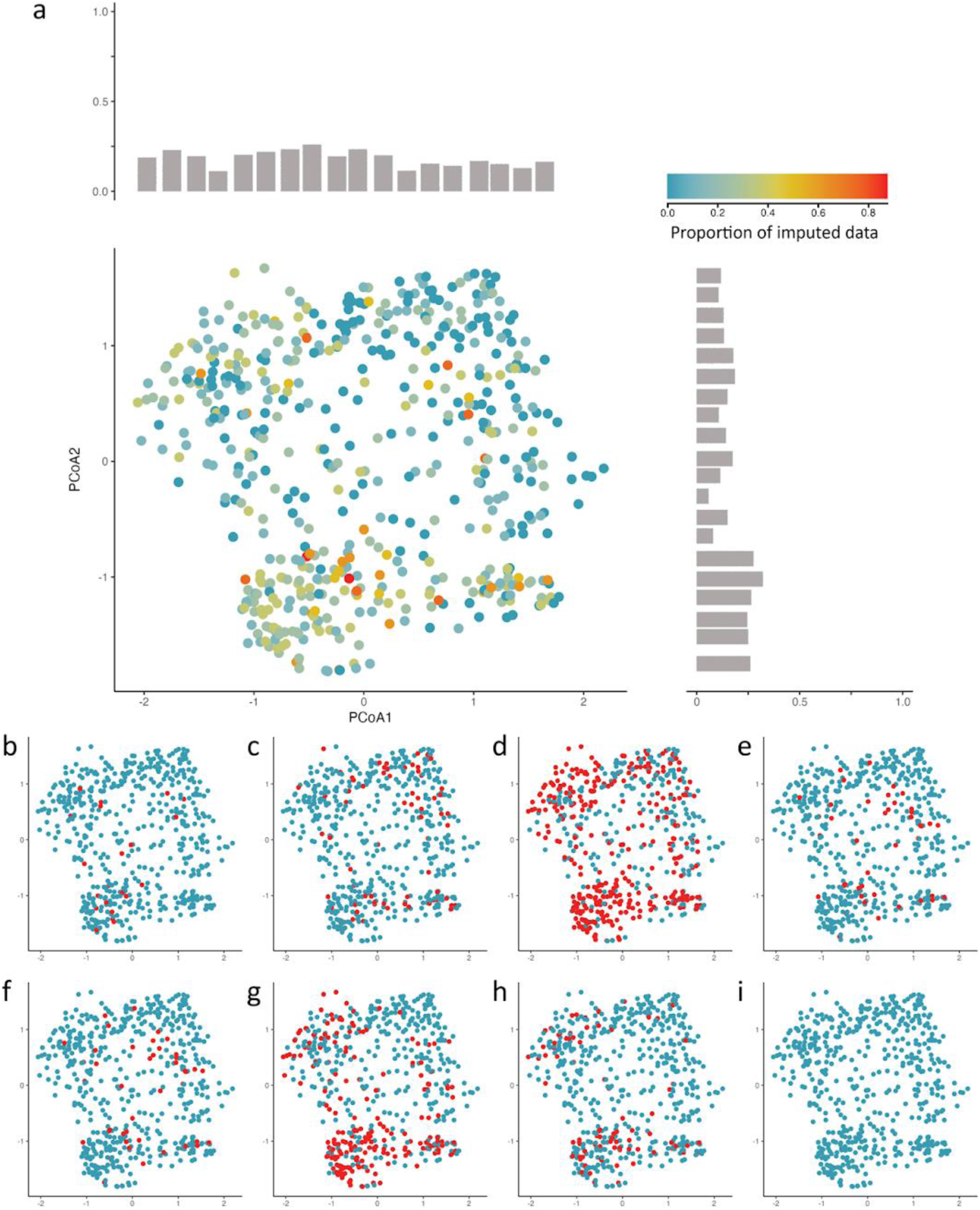
| The distribution of imputed values across the shark trait space. (a) Distribution of the proportion of imputed data across species, projected onto the first two principal coordinate (PCoA) axes. Points are coloured according to the proportion of imputed data (blue = low, red = high). Grey bars show marginal distributions of imputation density across species. Panels (b–i) show locations in trait space where imputation occurred for individual traits: (b) body length, (c) maximum depth, (d) trophic level, (e) body shape variation PC1, (f) body shape variation PC2, (g) growth ratio, (h) reproductive guild, and (i) habitat preference. Each red point indicates a species with imputed values for the given trait, while blue points represent species with observed values.

**Fig. S4.**
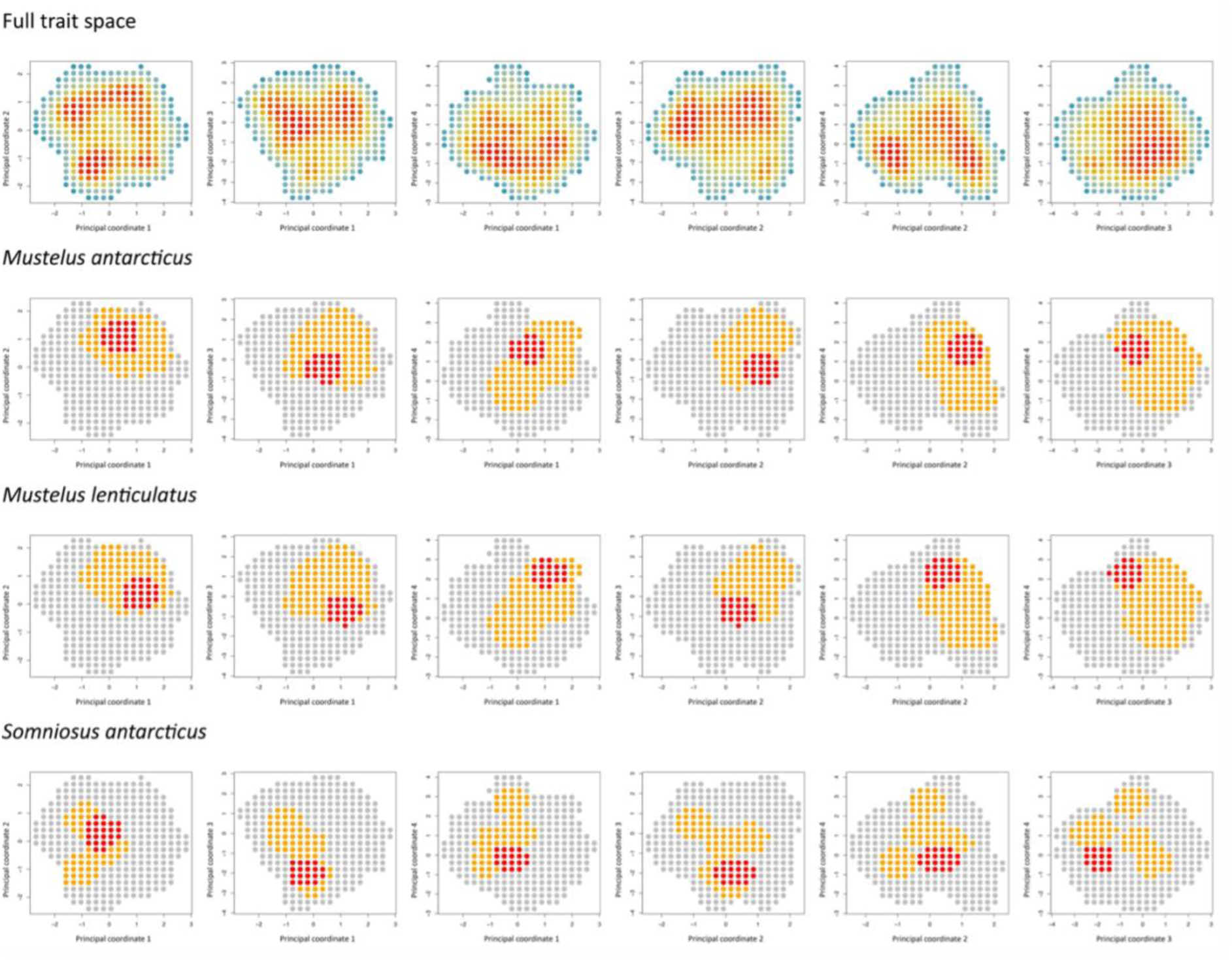
| Occupation of trait space for selected species and their congeners. Probability of occupancy of full trait space (a). Area of trait space occupied by all species in Mustelus genus (orange) and space occupied by Mustelus antarcticus (b) and Mustelus lenticulatus (c). Area of trait space occupied by all species in Somniosus genus (orange) and space occupied by Somniosus antarcticus (red; d).

**Fig. S5.**
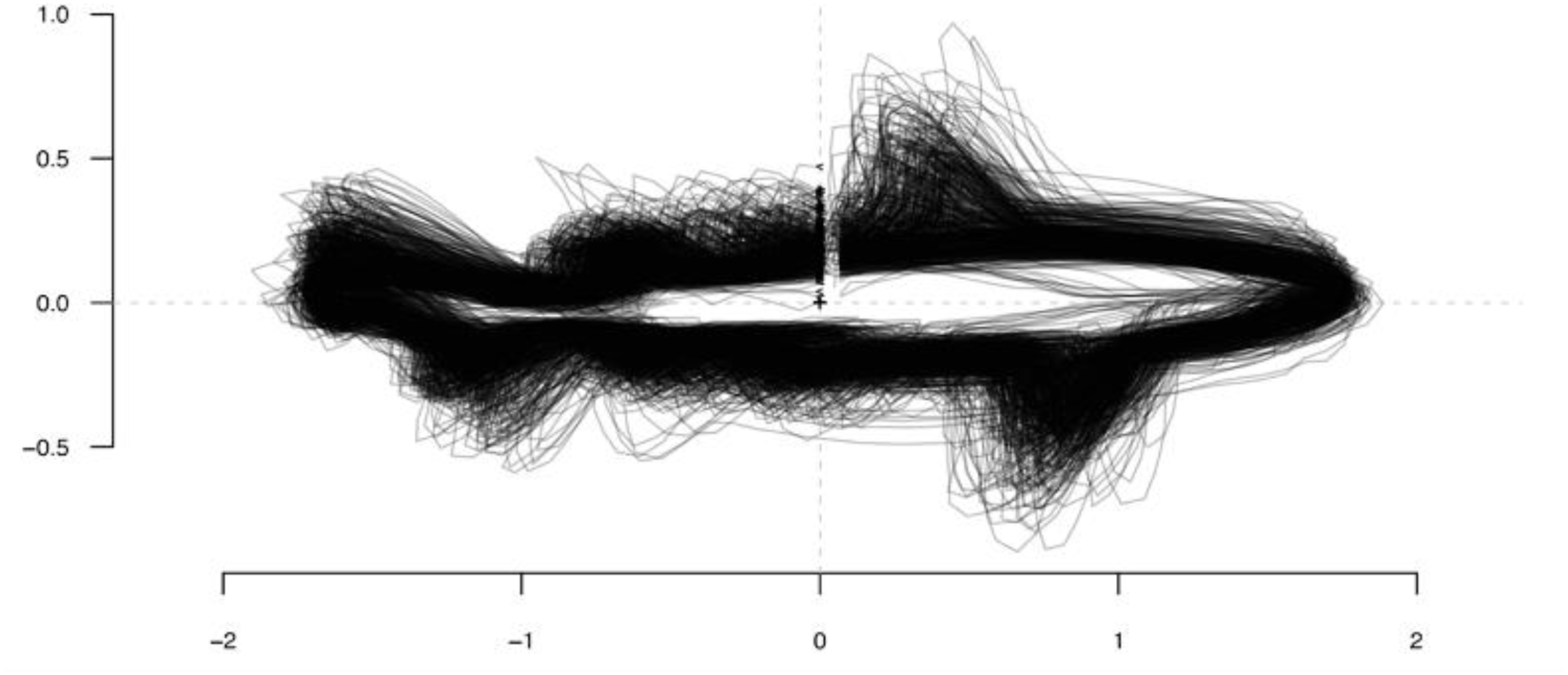
| Stacked shark outlines. Scaled and centred outlines to show the body shape silhouettes of 515 images of shark species collected from Sharks of the World: A Complete Guide^11^.

**Fig. S6.**
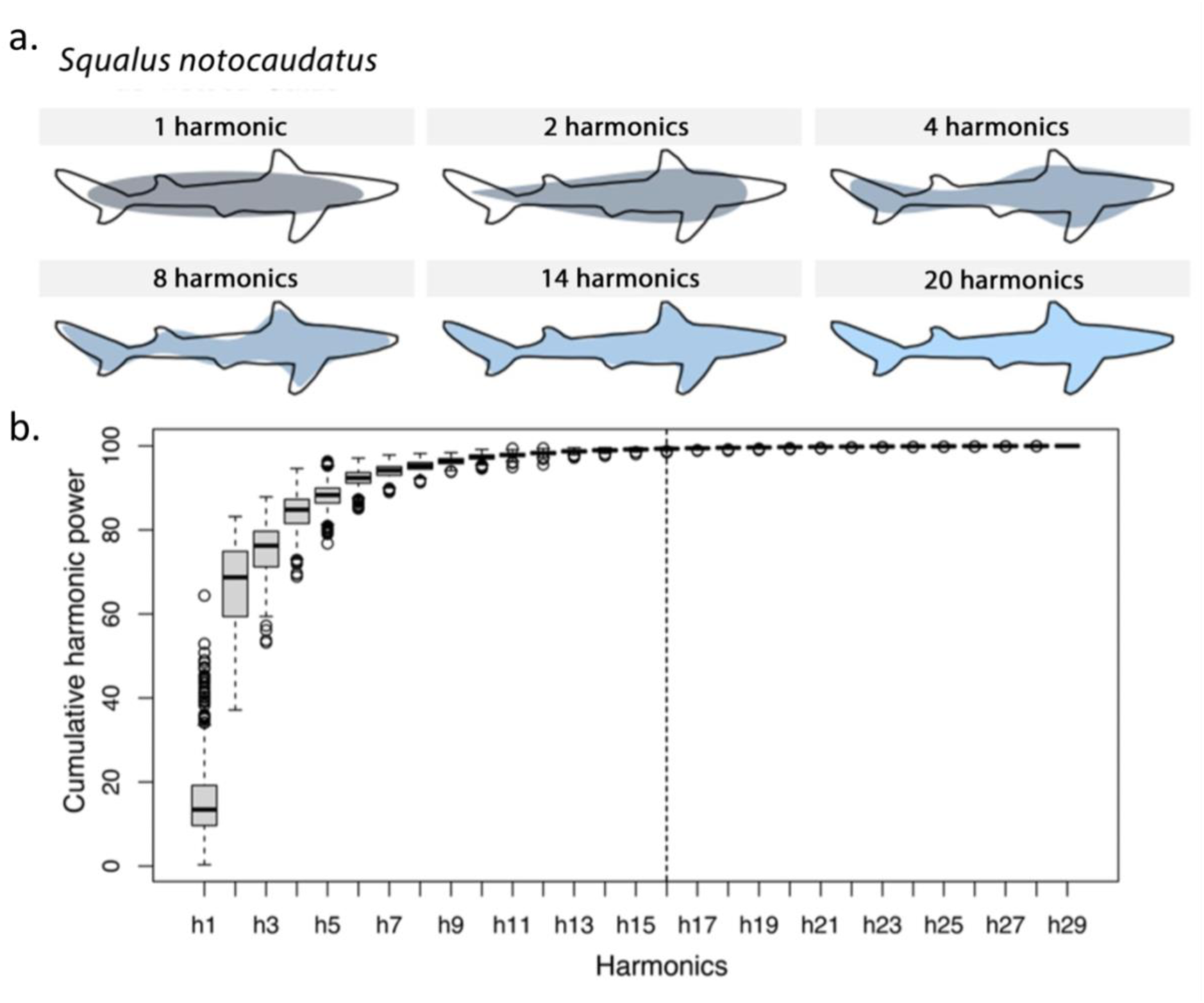
| Harmonic power required to fit shark outline. a) Number of harmonics required to gather cumulative harmonic power from the outlines of sharks following elliptical fourier analysis. b) Boxplots to show the harmonic power for the sum of harmonics (h1-h29) for the body shape elliptical fourier analysis. 16 harmonics were selected to gather 99% of cumulated harmonic power.

**Fig. S7.**
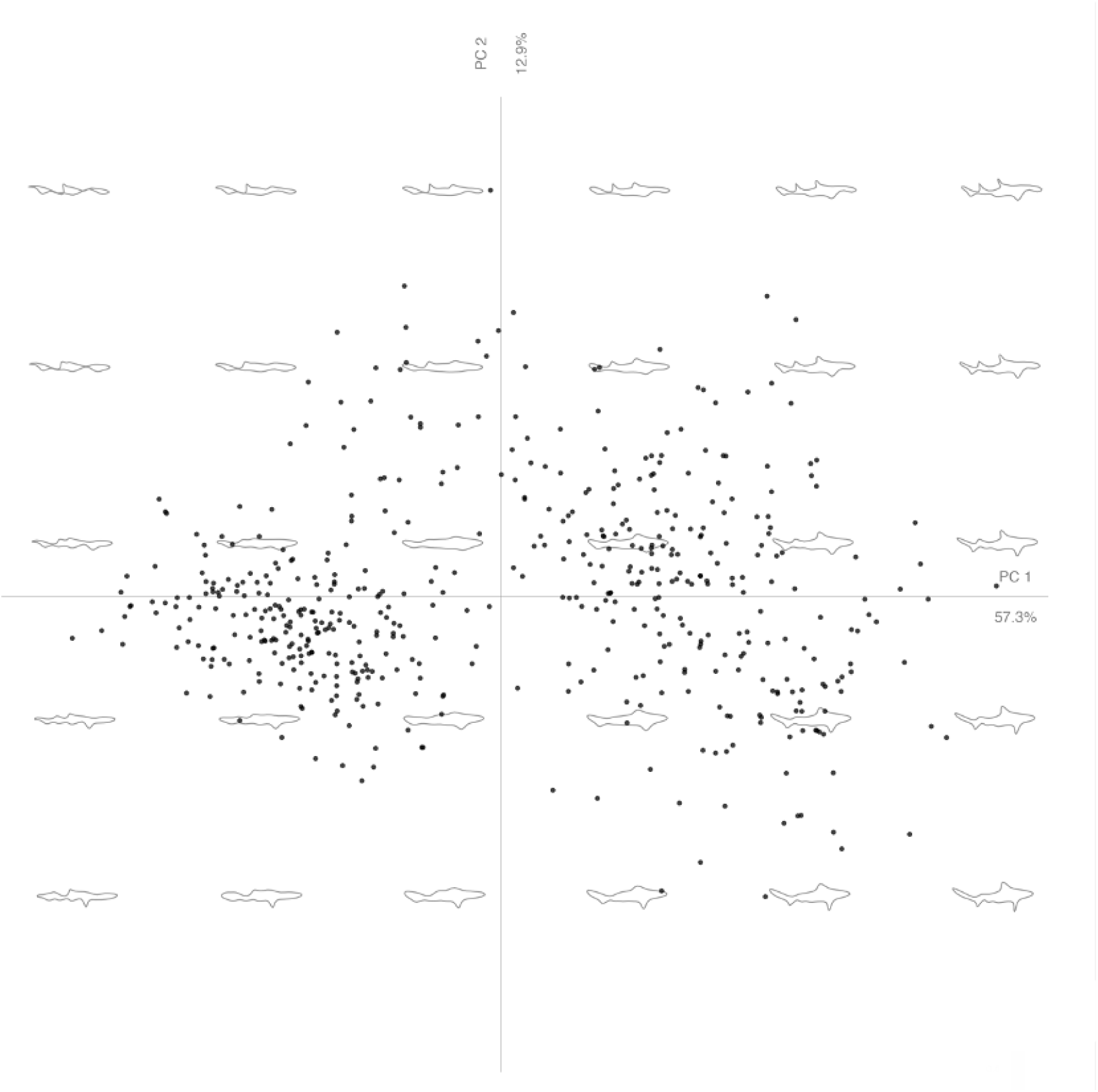
| Principal coordinate analysis of shark body shape elliptical fourier analysis. The first and second principal component from a principal component analysis of the body shape elliptical fourier analysis. Coordinates quantify variation in shape of shark species, with outlines representing mean body shape for areas shown on the graph.

**Table S1.**
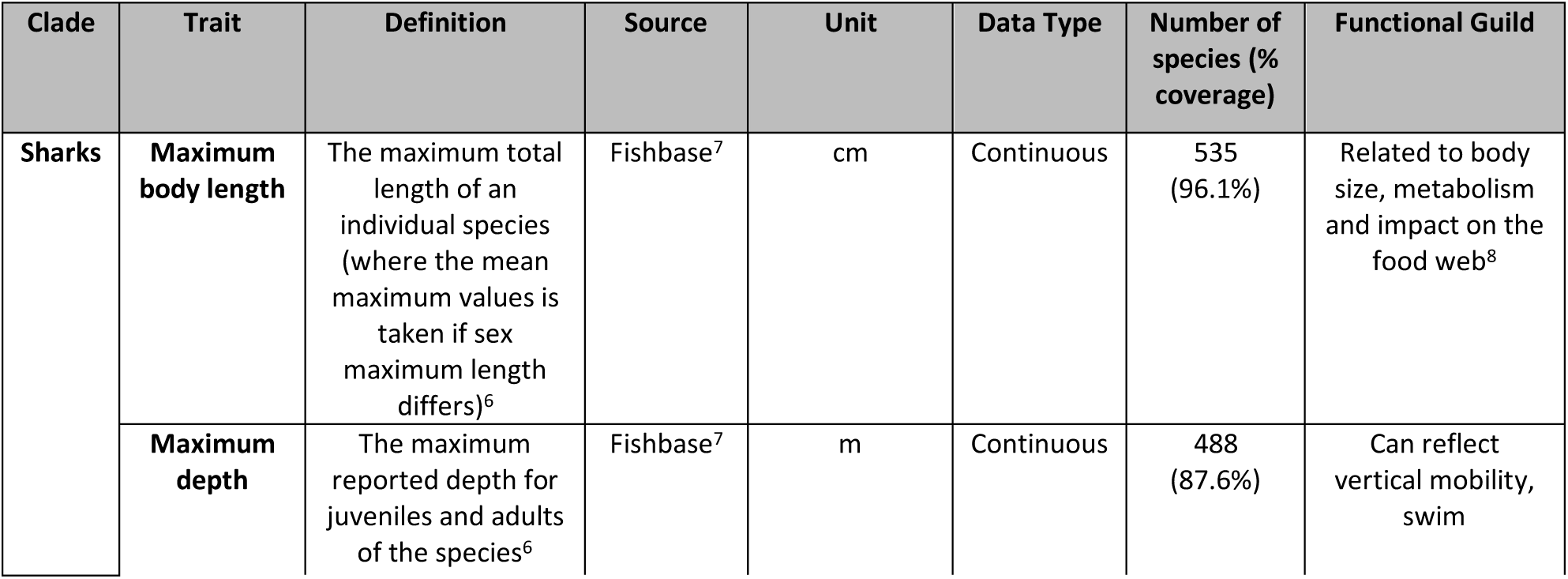
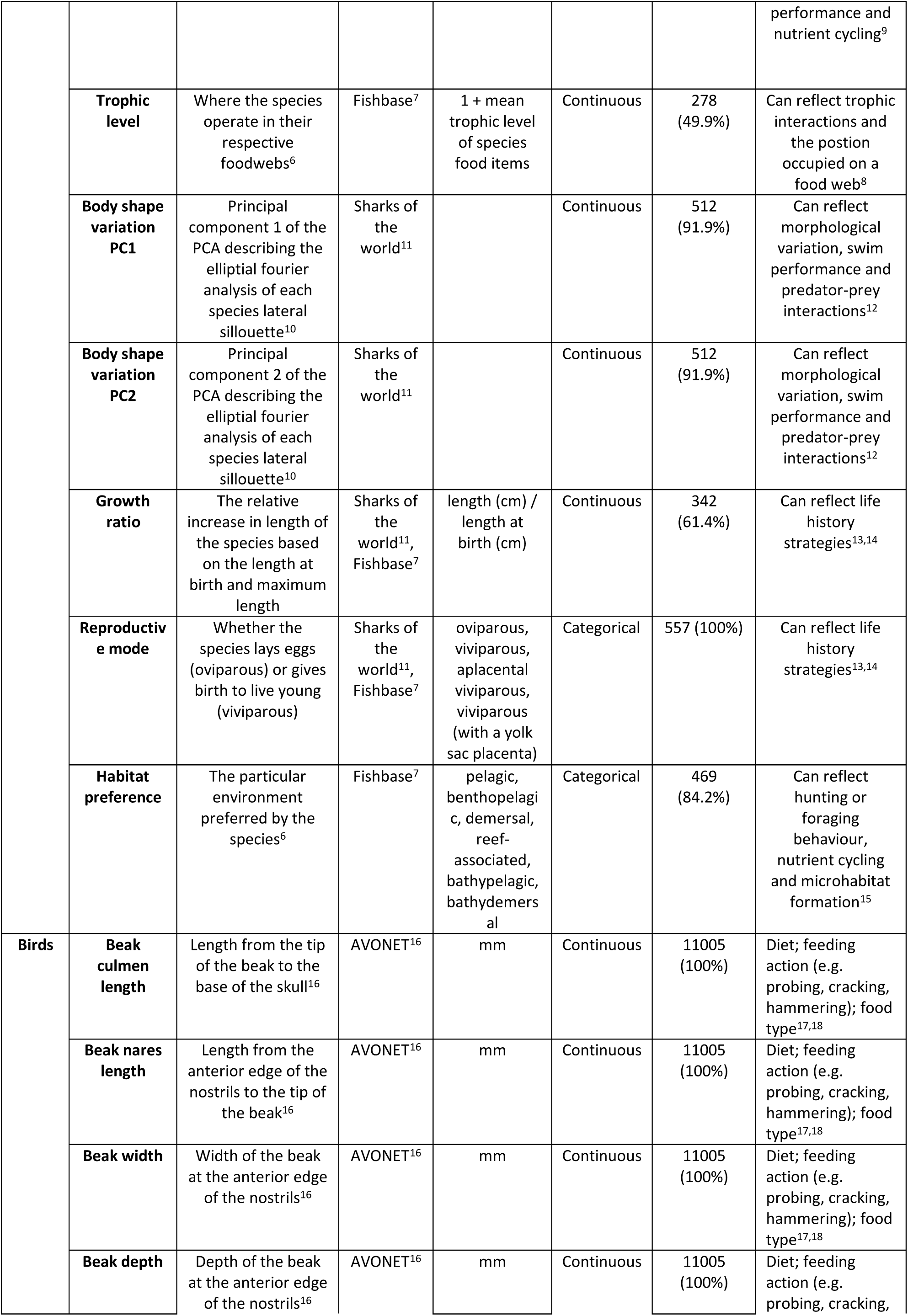
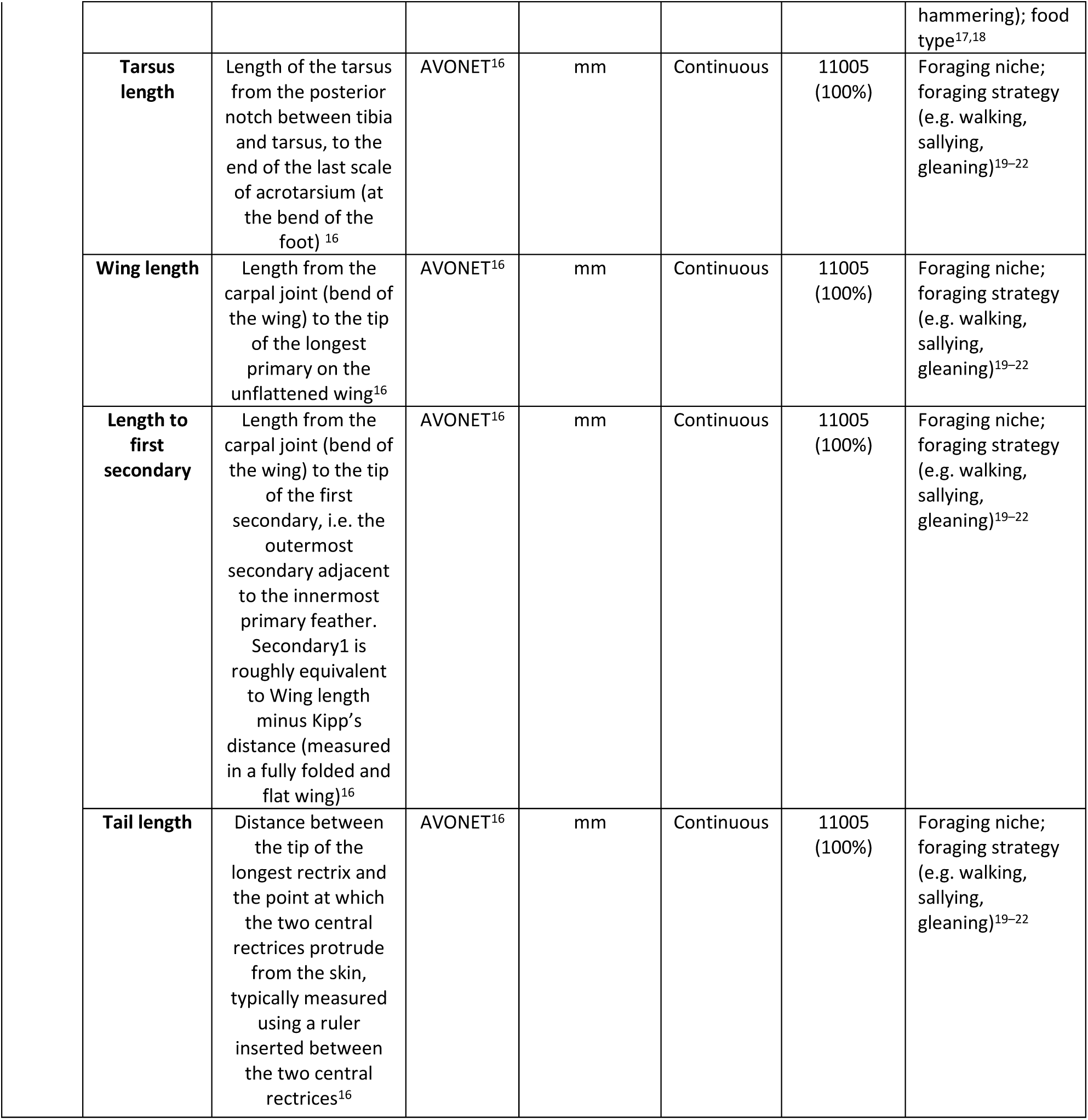
| Trait data. Traits used to build trait space and the functional guild each trait represents for both birds and sharks.

| Clade | Trait | Definition | Source | Unit | Data Type | Number of species (% coverage) | Functional Guild |
| --- | --- | --- | --- | --- | --- | --- | --- |
| Sharks | Maximum body length | The maximum total length of an individual species (where the mean maximum values is taken if sex maximum length differs)<sup>6</sup> | Fishbase<sup>7</sup> | cm | Continuous | 535 (96.1%) | Related to body size, metabolism and impact on the food web<sup>8</sup> |
|  | Maximum depth | The maximum reported depth for juveniles and adults of the species<sup>6</sup> | Fishbase<sup>7</sup> | m | Continuous | 488 (87.6%) | Can reflect vertical mobility, swim performance and nutrient cycling<sup>9</sup> |
|  | **Trophic level** | Where the species operate in their respective foodwebs<sup>6</sup> | Fishbase<sup>7</sup> | 1 + mean trophic level of species food items | Continuous | 278 (49.9%) | Can reflect trophic interactions and the position occupied on a food web<sup>8</sup> |
|  | **Body shape variation PC1** | Principal component 1 of the PCA describing the elliptical fourier analysis of each species lateral silhouette<sup>10</sup> | Sharks of the world<sup>11</sup> |  | Continuous | 512 (91.9%) | Can reflect morphological variation, swim performance and predator-prey interactions<sup>12</sup> |
|  | **Body shape variation PC2** | Principal component 2 of the PCA describing the elliptical fourier analysis of each species lateral silhouette<sup>10</sup> | Sharks of the world<sup>11</sup> |  | Continuous | 512 (91.9%) | Can reflect morphological variation, swim performance and predator-prey interactions<sup>12</sup> |
|  | **Growth ratio** | The relative increase in length of the species based on the length at birth and maximum length | Sharks of the world<sup>11</sup>, Fishbase<sup>7</sup> | length (cm) / length at birth (cm) | Continuous | 342 (61.4%) | Can reflect life history strategies<sup>13,14</sup> |
|  | **Reproductive mode** | Whether the species lays eggs (oviparous) or gives birth to live young (viviparous) | Sharks of the world<sup>11</sup>, Fishbase<sup>7</sup> | oviparous, viviparous, aplacental viviparous, viviparous (with a yolk sac placenta) | Categorical | 557 (100%) | Can reflect life history strategies<sup>13,14</sup> |
|  | **Habitat preference** | The particular environment preferred by the species<sup>6</sup> | Fishbase<sup>7</sup> | pelagic, benthopelagic, demersal, reef-associated, bathypelagic, bathydemersal | Categorical | 469 (84.2%) | Can reflect hunting or foraging behaviour, nutrient cycling and microhabitat formation<sup>15</sup> |
| **Birds** | **Beak culmen length** | Length from the tip of the beak to the base of the skull<sup>16</sup> | AVONET<sup>16</sup> | mm | Continuous | 11005 (100%) | Diet; feeding action (e.g. probing, cracking, hammering); food type<sup>17,18</sup> |
|  | **Beak nares length** | Length from the anterior edge of the nostrils to the tip of the beak<sup>16</sup> | AVONET<sup>16</sup> | mm | Continuous | 11005 (100%) | Diet; feeding action (e.g. probing, cracking, hammering); food type<sup>17,18</sup> |
|  | **Beak width** | Width of the beak at the anterior edge of the nostrils<sup>16</sup> | AVONET<sup>16</sup> | mm | Continuous | 11005 (100%) | Diet; feeding action (e.g. probing, cracking, hammering); food type<sup>17,18</sup> |
|  | **Beak depth** | Depth of the beak at the anterior edge of the nostrils<sup>16</sup> | AVONET<sup>16</sup> | mm | Continuous | 11005 (100%) | Diet; feeding action (e.g. probing, cracking, |

|  |  |  |  |  |  |  |
| --- | --- | --- | --- | --- | --- | --- |
|  |  |  |  |  |  | hammering); food type<sup>17,18</sup> |
| **Tarsus length** | Length of the tarsus from the posterior notch between tibia and tarsus, to the end of the last scale of acrotarsium (at the bend of the foot)<sup>16</sup> | AVONET<sup>16</sup> | mm | Continuous | 11005 (100%) | Foraging niche; foraging strategy (e.g. walking, sallying, gleaning)<sup>19–22</sup> |
| **Wing length** | Length from the carpal joint (bend of the wing) to the tip of the longest primary on the unflattened wing<sup>16</sup> | AVONET<sup>16</sup> | mm | Continuous | 11005 (100%) | Foraging niche; foraging strategy (e.g. walking, sallying, gleaning)<sup>19–22</sup> |
| **Length to first secondary** | Length from the carpal joint (bend of the wing) to the tip of the first secondary, i.e. the outermost secondary adjacent to the innermost primary feather. Secondary1 is roughly equivalent to Wing length minus Kipp's distance (measured in a fully folded and flat wing)<sup>16</sup> | AVONET<sup>16</sup> | mm | Continuous | 11005 (100%) | Foraging niche; foraging strategy (e.g. walking, sallying, gleaning)<sup>19–22</sup> |
| **Tail length** | Distance between the tip of the longest rectrix and the point at which the two central rectrices protrude from the skin, typically measured using a ruler inserted between the two central rectrices<sup>16</sup> | AVONET<sup>16</sup> | mm | Continuous | 11005 (100%) | Foraging niche; foraging strategy (e.g. walking, sallying, gleaning)<sup>19–22</sup> |

**Table S2.**
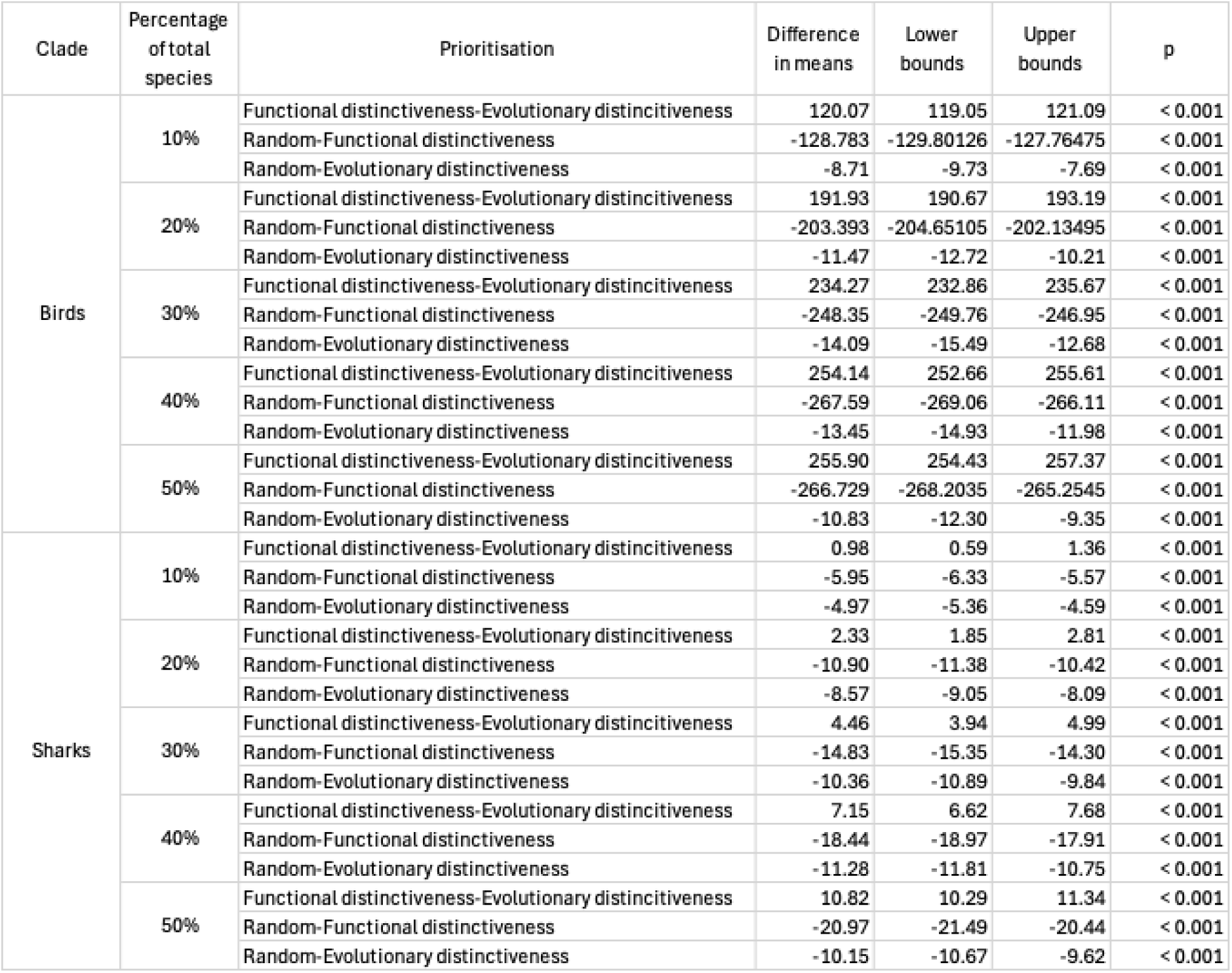
| ANOVA pairwise comparisons between prioritisation strategies at capturing intentionally exploited bird and shark species with Tukey’s Honestly Significant Difference post hoc test.

## List of Supplementary data

Data S1 – Bird indices

Data S2 – Shark indices

Data S3 – Trait database for birds

Data S4 – Trait database for sharks

Data S5 – PCA shape data

## References

1. Mace, G. M., Norris, K. & Fitter, A. H. Biodiversity and ecosystem services: a multilayered relationship. Trends in Ecology & Evolution 27, 19–26 (2012).

2. IUCN Red List of Threatened Species. The IUCN Red List of Threatened Species. IUCN Red List of Threatened Species https://www.iucnredlist.org/en (2025).

3. Hoekstra, J. M., Boucher, T. M., Ricketts, T. H. & Roberts, C. Confronting a biome crisis: global disparities of habitat loss and protection. Ecology Letters 8, 23–29 (2005).

4. Mittermeier, R. A. et al. Wilderness and biodiversity conservation. Proceedings of the National Academy of Sciences 100, 10309–10313 (2003).

5. IPCC. Global Warming of 1.5°C: IPCC Special Report on Impacts of Global Warming of 1.5°C above Pre-Industrial Levels in Context of Strengthening Response to Climate Change, Sustainable Development, and Efforts to Eradicate Poverty. (Cambridge University Press, Cambridge, 2022).

6. Foden, W. B. et al. Climate change vulnerability assessment of species. WIREs Climate Change 10, e551 (2019).

7. Isaac, N. J. B., Turvey, S. T., Collen, B., Waterman, C. & Baillie, J. E. M. Mammals on the EDGE: Conservation Priorities Based on Threat and Phylogeny. PLOS ONE 2, e296 (2007).

8. Cadotte, M. W., Carscadden, K. & Mirotchnick, N. Beyond species: functional diversity and the maintenance of ecological processes and services. Journal of Applied Ecology 48, 1079–1087 (2011).

9. Violle, C. et al. Let the concept of trait be functional! Oikos 116, 882–892 (2007).

10. Palacio, F. X. et al. A protocol for reproducible functional diversity analyses. Ecography 2022, e06287 (2022).

11. Miatta, M., Bates, A. E. & Snelgrove, P. V. R. Incorporating biological traits into conservation strategies. Annu. Rev. Mar. Sci. 13, 421–443 (2021).

12. Convention on Biological Diversity. Kunming-Montreal Global Biodiversity Framework. Tech. rep. Montreal, Canada (2022).

13. Tobias, J. A., Bullock, J. M., Dicks, L. V., Forester, B. R. & Razgour, O. Biodiversity conservation requires integration of species-centric and process-based strategies. Proceedings of the National Academy of Sciences 122, e2410936122 (2025).

14. de Bello, F., Botta-Dukát, Z., Lepš, J. & Fibich, P. Towards a more balanced combination of multiple traits when computing functional differences between species. Methods in Ecology and Evolution 12, 443–448 (2021).

15. Ali, J. R., Blonder, B. W., Pigot, A. L. & Tobias, J. A. Bird extinctions threaten to cause disproportionate reductions of functional diversity and uniqueness. Functional Ecology 37, 162–175 (2023).

16. Carmona, C. P. et al. Erosion of global functional diversity across the tree of life. Science Advances 7, eabf2675 (2021).

17. Oliveira, B. F., Scheffers, B. R. & Costa, G. C. Decoupled erosion of amphibians’ phylogenetic and functional diversity due to extinction. Global Ecology and Biogeography 29, 309–319 (2020).

18. Pollock, L. J., Thuiller, W. & Jetz, W. Large conservation gains possible for global biodiversity facets. Nature 546, 141–144 (2017).

19. Faith, D. P. Conservation evaluation and phylogenetic diversity. Biological Conservation 61, 1–10 (1992).

20. Gumbs, R. et al. The EDGE2 protocol: Advancing the prioritisation of Evolutionarily Distinct and Globally Endangered species for practical conservation action. PLOS Biology 21, e3001991 (2023).

21. Faith, D. P. Threatened Species and the Potential Loss of Phylogenetic Diversity: Conservation Scenarios Based on Estimated Extinction Probabilities and Phylogenetic Risk Analysis. Conservation Biology 22, 1461–1470 (2008).

22. Jensen, E. L., Mooers, A. Ø., Caccone, A. & Russello, M. A. I-HEDGE: determining the optimum complementary sets of taxa for conservation using evolutionary isolation. PeerJ 4, e2350 (2016).

23. Robuchon, M. et al. Revisiting species and areas of interest for conserving global mammalian phylogenetic diversity. Nat Commun 12, 3694 (2021).

24. Steel, M., Mimoto, A. & Mooers, A. Ø. Hedging Our Bets: The Expected Contribution of Species to Future Phylogenetic Diversity. Evol Bioinform Online 3, 117693430700300024 (2007).

25. Gouhier, T. C. & Pillai, P. Avoiding Conceptual and Mathematical Pitfalls When Developing Indices to Inform Conservation. Frontiers in Ecology and Evolution 8, (2020).

26. Pavoine, S. & Ricotta, C. Identifying functionally distinctive and threatened species. Biological Conservation 284, 110170 (2023).

27. Hidasi-Neto, J., Loyola, R. & Cianciaruso, M. V. Global and local evolutionary and ecological distinctiveness of terrestrial mammals: identifying priorities across scales. Diversity and Distributions 21, 548–559 (2015).

28. Pimiento, C. et al. Functional diversity of marine megafauna in the Anthropocene. Science Advances 6, eaay7650 (2020).

29. Mouillot, D. et al. Rare Species Support Vulnerable Functions in High-Diversity Ecosystems. PLOS Biology 11, e1001569 (2013).

30. Mammola, S., Carmona, C. P., Guillerme, T. & Cardoso, P. Concepts and applications in functional diversity. Functional Ecology 35, 1869–1885 (2021).

31. Tucker, C. M. et al. A guide to phylogenetic metrics for conservation, community ecology and macroecology. Biological Reviews 92, 698–715 (2017).

32. Pimiento, C. et al. Functional diversity of sharks and rays is highly vulnerable and supported by unique species and locations worldwide. Nat Commun 14, 7691 (2023).

33. Tobias, J. A. et al. AVONET: morphological, ecological and geographical data for all birds. Ecology Letters 25, 581–597 (2022).

34. Carmona, C. P., de Bello, F., Mason, N. W. H. & Lepš, J. Trait probability density (TPD): measuring functional diversity across scales based on TPD with R. Ecology 100, e02876 (2019).

35. Fraixedas, S. et al. A state-of-the-art review on birds as indicators of biodiversity: Advances, challenges, and future directions. Ecological Indicators 118, 106728 (2020).

36. Hughes, A. C. et al. Sampling biases shape our view of the natural world. Ecography 44, 1259–1269 (2021).

37. Pigot, A. L. et al. Macroevolutionary convergence connects morphological form to ecological function in birds. Nat Ecol Evol 4, 230–239 (2020).

38. Carmona, C. P., de Bello, F., Mason, N. W. H. & Lepš, J. Traits Without Borders: Integrating Functional Diversity Across Scales. Trends in Ecology & Evolution 31, 382– 394 (2016).

39. Hughes, E. C., Edwards, D. P. & Thomas, G. H. The homogenization of avian morphological and phylogenetic diversity under the global extinction crisis. Current Biology 32, 3830–3837.e3 (2022).

40. Matthews, T. J. et al. Threatened and extinct island endemic birds of the world: Distribution, threats and functional diversity. Journal of Biogeography 49, 1920–1940 (2022).

41. Gumbs, R. et al. Global conservation status of the jawed vertebrate Tree of Life. Nat Commun 15, 1101 (2024).

42. BirdLife International. Apteryx rowi. The IUCN Red List of Threatened Species 2021: e.T22732871A180757495. 10.2305/IUCN.UK.2021-3.RLTS.T22732871A180757495.en (2021).

43. Díaz, S. et al. Set ambitious goals for biodiversity and sustainability. Science 370, 411– 413 (2020).

44. Mace, G. M. et al. Approaches to defining a planetary boundary for biodiversity. Global Environmental Change 28, 289–297 (2014).

45. McKinney, M. L. High Rates of Extinction and Threat in Poorly Studied Taxa. Conservation Biology 13, 1273–1281 (1999).

46. Bland, L. M., Collen, B., Orme, C. D. L. & Bielby, J. Predicting the conservation status of data-deficient species. Conserv Biol 29, 250–259 (2015).

47. Bachman, S. P., Brown, M. J. M., Leão, T. C. C., Nic Lughadha, E. & Walker, B. E. Extinction risk predictions for the world’s flowering plants to support their conservation. New Phytologist 242, 797–808 (2024).

48. Weedop, K. B., Mooers, A. Ø., Tucker, C. M. & Pearse, W. D. The effect of phylogenetic uncertainty and imputation on EDGE Scores. Animal Conservation 22, 527–536 (2019).

49. Stewart, K. et al. Functional diversity metrics can perform well with highly incomplete data sets. Methods in Ecology and Evolution 14, 2856–2872 (2023).

50. Dulvy, N. K. et al. Overfishing drives over one-third of all sharks and rays toward a global extinction crisis. Current Biology 31, 4773–4787.e8 (2021).

51. Stekhoven, D. J. & Bühlmann, P. MissForest—non-parametric missing value imputation for mixed-type data. Bioinformatics 28, 112–118 (2012).

52. Gendre, M., Hauffe, T., Pimiento, C. & Silvestro, D. Benchmarking imputation methods for categorical biological data. Methods in Ecology and Evolution 15, 1624–1638 (2024).

53. May, J. A., Feng, Z. & Adamowicz, S. J. A real data-driven simulation strategy to select an imputation method for mixed-type trait data. PLOS Computational Biology 19, e1010154 (2023).

54. White, W. T., Kyne, P. M. & Harris, M. Lost before found: A new species of whaler shark Carcharhinus obsolerus from the Western Central Pacific known only from historic records. PLOS ONE 14, e0209387 (2019).

55. Violle, C. et al. Functional Rarity: The Ecology of Outliers. Trends in Ecology & Evolution 32, 356–367 (2017).

56. Cadotte, M. W. Functional traits explain ecosystem function through opposing mechanisms. Ecology Letters 20, 989–996 (2017).

57. Díaz, S. et al. Functional traits, the phylogeny of function, and ecosystem service vulnerability. Ecology and Evolution 3, 2958–2975 (2013).

58. CBD. Monitoring framework for the Kunming-Montreal global biodiversity framework. in Conference of the parties to the convention on biological diversity fifteenth meeting. p. CBD/COP/DEC/15/5 (2022).

59. Mazel, F. et al. Prioritizing phylogenetic diversity captures functional diversity unreliably. Nat Commun 9, 2888 (2018).

60. Cadotte, M., Albert, C. H. & Walker, S. C. The ecology of differences: assessing community assembly with trait and evolutionary distances. Ecology Letters 16, 1234– 1244 (2013).

61. Cardoso, P. et al. Calculating functional diversity metrics using neighbor-joining trees. Ecography 2024, e07156 (2024).

62. Matthews, T. J. et al. The global loss of avian functional and phylogenetic diversity from anthropogenic extinctions. Science 386, 55–60 (2024).

63. Kattge, J. et al. TRY plant trait database–enhanced coverage and open access. Global change biology 26, 119–188 (2020).

64. Froese, R. & Pauly, D. FishBase. https://www.fishbase.org (2023).

65. Meiri, S. SquamBase—A database of squamate (Reptilia: Squamata) traits. Global Ecology and Biogeography 33, e13812 (2024).

66. Jones, K. E. et al. PanTHERIA: a species-level database of life history, ecology, and geography of extant and recently extinct mammals: Ecological Archives E090-184. Ecology 90, 2648–2648 (2009).

67. Buuren, S. V. & Groothuis-Oudshoorn, K. **mice** : Multivariate Imputation by Chained Equations in R. J. Stat. Soft. 45, (2011).

68. Goolsby, E. W., Bruggeman, J. & Ané, C. Rphylopars: fast multivariate phylogenetic comparative methods for missing data and within-species variation. Methods in Ecology and Evolution 8, 22–27 (2017).

69. Blomberg, S. P. & Todorov, O. S. The fallacy of single imputation for trait databases: Use multiple imputation instead. Methods in Ecology and Evolution 16, 658–667 (2025).

70. Oskyrko, O., Mi, C., Meiri, S. & Du, W. ReptTraits: a comprehensive dataset of ecological traits in reptiles. Sci Data 11, 243 (2024).

71. Oliveira, B. F., São-Pedro, V. A., Santos-Barrera, G., Penone, C. & Costa, G. C. AmphiBIO, a global database for amphibian ecological traits. Sci Data 4, 170123 (2017).

72. Mull, C. G. et al. Sharkipedia: a curated open access database of shark and ray life history traits and abundance time-series. Sci Data 9, 559 (2022).

73. Ahnelt, H., Sauberer, M., Ramler, D., Koch, L. & Pogoreutz, C. Negative allometric growth during ontogeny in the large pelagic filter-feeding basking shark. Zoomorphology 139, 71–83 (2020).

74. Curnick, D. J., Carlisle, A. B., Gollock, M. J., Schallert, R. J. & Hussey, N. E. Evidence for dynamic resource partitioning between two sympatric reef shark species within the British Indian Ocean Territory. Journal of Fish Biology 94, 680–685 (2019).

75. Vane-Wright, R. I., Humphries, C. J. & Williams, P. H. What to protect?—Systematics and the agony of choice. Biological Conservation 55, 235–254 (1991).

76. Collen, B. et al. Investing in evolutionary history: implementing a phylogenetic approach for mammal conservation. Philosophical Transactions of the Royal Society B: Biological Sciences 366, 2611–2622 (2011).

77. Forest, F. et al. Preserving the evolutionary potential of floras in biodiversity hotspots. Nature 445, 757–760 (2007).

78. Gumbs, R. et al. Conserving avian evolutionary history can effectively safeguard future benefits for people. Science Advances 9, eadh4686 (2023).

79. Molina-Venegas, R. Conserving evolutionarily distinct species is critical to safeguard human well-being. Sci Rep 11, 24187 (2021).

80. Rosindell, J., Manson, K., Gumbs, R., Pearse, W. D. & Steel, M. Phylogenetic Biodiversity Metrics Should Account for Both Accumulation and Attrition of Evolutionary Heritage. Systematic Biology 73, 158–182 (2024).

81. Kemp, T. S. The Origin of Higher Taxa: Palaeobiological, Developmental, and Ecological Perspectives. (University of Chicago Press, Chicago, IL, 2015).

82. Tucker, C. M., Davies, T. J., Cadotte, M. W. & Pearse, W. D. On the relationship between phylogenetic diversity and trait diversity. Ecology 99, 1473–1479 (2018).

83. Chichorro, F. et al. Trait-based prediction of extinction risk across terrestrial taxa. Biological Conservation 274, 109738 (2022).

84. Stewart, K. et al. Threat reduction must be coupled with targeted recovery programmes to conserve global bird diversity. Nat Ecol Evol 1–13 (2025) doi:10.1038/s41559-025-02746-z.

85. Molina-Venegas, R., Rodríguez, M. Á., Pardo-de-Santayana, M., Ronquillo, C. & Mabberley, D. J. Maximum levels of global phylogenetic diversity efficiently capture plant services for humankind. Nat Ecol Evol 5, 583–588 (2021).

86. Ripple, W. J. et al. Extinction risk is most acute for the world’s largest and smallest vertebrates. Proceedings of the National Academy of Sciences 114, 10678–10683 (2017).

87. Ripple, W. J. et al. Are we eating the world’s megafauna to extinction? Conservation Letters 12, e12627 (2019).

88. Toomes, A. et al. Drivers of the Australian native pet trade: The role of species traits, socioeconomic attributes and regulatory systems. Journal of Applied Ecology 59, 1268– 1278 (2022).

89. Branch, T. A., Lobo, A. S. & Purcell, S. W. Opportunistic exploitation: an overlooked pathway to extinction. Trends in Ecology & Evolution 28, 409–413 (2013).

90. Blowes, S. A. et al. The geography of biodiversity change in marine and terrestrial assemblages. Science 366, 339–345 (2019).

91. Hillebrand, H. et al. Biodiversity change is uncoupled from species richness trends: Consequences for conservation and monitoring. Journal of Applied Ecology 55, 169–184 (2018).

92. Cooke, R. S. C., Eigenbrod, F. & Bates, A. E. Projected losses of global mammal and bird ecological strategies. Nat Commun 10, 2279 (2019).

93. IPBES. Global Assessment Report on Biodiversity and Ecosystem Services of the Intergovernmental Science-Policy Platform on Biodiversity and Ecosystem Services. https://zenodo.org/records/6417333 (2019) doi:10.5281/zenodo.6417333.

94. IUCN. IUCN Red List Categories and Criteria, Version 3.1. (IUCN Species Survival Commission, 2001).

95. BirdLife International. BirdLife Taxonomic Checklist v6. (2021).

96. R Core Team. R: A Language and Environment for Statistical Computing. R Foundation for Statistical Computing (2023).

97. Jetz, W. et al. Global Distribution and Conservation of Evolutionary Distinctness in Birds. Current Biology 24, 919–930 (2014).

98. Safi, K., Armour-Marshall, K., Baillie, J. E. M. & Isaac, N. J. B. Global Patterns of Evolutionary Distinct and Globally Endangered Amphibians and Mammals. PLOS ONE 8, e63582 (2013).

99. McClure, C. J. W. et al. Conserving the evolutionary history of birds. Conservation Biology 37, e14141 (2023).

100. BirdLife International. BirdLife Taxonomic Checklist. (2024).

101. BirdLife International. The World Database of Key Biodiversity Areas. (2025).

102. Lansley, T. P., Crowe, O., Butchart, S. H. M., Edwards, D. P. & Thomas, G. H. Effectiveness of Key Biodiversity Areas in representing global avian diversity. Conservation Biology n/a, e70000 (2025).

103. Brooks, T. M. et al. Measuring Terrestrial Area of Habitat (AOH) and Its Utility for the IUCN Red List. Trends in Ecology & Evolution 34, 977–986 (2019).

104. IUCN Red List of Threatened Species. The IUCN Red List of Threatened Species. IUCN Red List of Threatened Species https://www.iucnredlist.org/en (2023).

105. Stein, R. W. et al. Global priorities for conserving the evolutionary history of sharks, rays and chimaeras. Nat Ecol Evol 2, 288–298 (2018).

106. Ebert, D. A., Dando, M. & Fowler, S. Sharks of the World: A Complete Guide. (Princeton University Press, 2021).

107. Villéger, S., Brosse, S., Mouchet, M., Mouillot, D. & Vanni, M. J. Functional ecology of fish: current approaches and future challenges. Aquat Sci 79, 783–801 (2017).

108. Siders, Z. A. et al. Functional and phylogenetic diversity of sharks in the Northeastern Pacific. Journal of Biogeography **n/a**, (2022).

109. Henderson, C. J. et al. Long term declines in the functional diversity of sharks in the coastal oceans of eastern Australia. Commun Biol 7, 1–8 (2024).

110. Adobe Inc. Adobe Photoshop. (2023).

111. Bonhomme, V., Picq, S., Gaucherel, C. & Claude, J. Momocs: Outline Analysis Using R. Journal of Statistical Software 56, 1–24 (2014).

112. Dray, S. & Dufour, A.-B. The ade4 Package: Implementing the Duality Diagram for Ecologists. Journal of Statistical Software 22, 1–20 (2007).

## References

1. Pigot, A. L. et al. Macroevolutionary convergence connects morphological form to ecological function in birds. Nat Ecol Evol 4, 230–239 (2020).

2. Williams, P. J. & Curley, S. R. Functional and phylogenetic convergence of winter and breeding bird communities in the northeastern US. Ecography e07717 (2025) doi:10.1002/ecog.07717.

3. Grenié, M., Denelle, P., Tucker, C. M., Munoz, F. & Violle, C. funrar: An R package to characterize functional rarity. Diversity and Distributions 23, 1365–1371 (2017).

4. Carmona, C. P., de Bello, F., Mason, N. W. H. & Lepš, J. Trait probability density (TPD): measuring functional diversity across scales based on TPD with R. Ecology 100, e02876 (2019).

5. IUCN Red List of Threatened Species. The IUCN Red List of Threatened Species. IUCN Red List of Threatened Species https://www.iucnredlist.org/en (2025).

6. Froese, R. & Pauly, D. FishBase 2000: Concepts, design and data sources. ICLARM (2000).

7. Froese, R. & Pauly, D. FishBase. https://www.fishbase.org (2023).

8. Henderson, C. J. et al. Long term declines in the functional diversity of sharks in the coastal oceans of eastern Australia. Commun Biol 7, 1–8 (2024).

9. Andrzejaczek, S. et al. Diving into the vertical dimension of elasmobranch movement ecology. Science Advances 8, eabo1754 (2022).

10. Siders, Z. A. et al. Functional and phylogenetic diversity of sharks in the Northeastern Pacific. Journal of Biogeography n/a, (2022).

11. Ebert, D. A., Dando, M. & Fowler, S. Sharks of the World: A Complete Guide. (Princeton University Press, 2021).

12. Sternes, P. C. & Shimada, K. Body forms in sharks (Chondrichthyes: Elasmobranchii) and their functional, ecological, and evolutionary implications. Zoology 140, 125799 (2020).

13. Cortés, E. Life history patterns, demography, and population dynamics. Biology of sharks and their relatives 449–469 (2004).

14. Cortés, E. Life History Patterns and Correlations in Sharks. Reviews in Fisheries Science 8, 299–344 (2000).

15. Sorenson, L., Santini, F. & Alfaro, M. E. The effect of habitat on modern shark diversification. Journal of Evolutionary Biology 27, 1536–1548 (2014).

16. Tobias, J. A. et al. AVONET: morphological, ecological and geographical data for all birds. Ecology Letters 25, 581–597 (2022).

17. Felice, R. N., Tobias, J. A., Pigot, A. L. & Goswami, A. Dietary niche and the evolution of cranial morphology in birds. Proceedings of the Royal Society B: Biological Sciences 286, 20182677 (2019).

18. Sayol, F., Reijenga, B. R., Tobias, J. A. & Pigot, A. L. Ecophysical constraints on avian adaptation and diversification. Current Biology 35, 1326–1336.e6 (2025).

19. Fitzpatrick. Tail length in birds in relation to tail shape, general flight ecology and sexual selection. Journal of Evolutionary Biology 12, 49–60 (1999).

20. Einoder, L. D. & Richardson, A. M. M. Aspects of the Hindlimb Morphology of Some Australian Birds of Prey: A Comparative and Quantitative Study. Auk 124, 773–788 (2007).

21. Miles, D. B. & Ricklefs, R. E. The Correlation Between Ecology and Morphology in Deciduous Forest Passerine Birds. Ecology 65, 1629–1640 (1984).

22. Miles, D. B., Ricklefs, R. E. & Travis, J. Concordance of Ecomorphological Relationships in Three Assemblages of Passerine Birds. The American Naturalist 129, 347–364 (1987).

